# Methylation-mediated retuning on the enhancer-to-silencer activity scale of networked regulatory elements guides driver-gene misregulation

**DOI:** 10.1101/2021.03.02.433521

**Authors:** Y. Edrei, R. levy, A. Marom, B. Radlwimmer, A. Hellman

**Author notes:** Correspondence to: Asaf Hellman < >.

## Abstract

Cancers arise when particular disease-driving genes adopt abnormal functions, but analyses of coding and regulatory sequences leave many of these abnormalities unexplained. We developed a strategy to explore alternations in the regulatory effects of silencers and enhancers in cancer tumors. Applying the method to 177 gene regulatory domains in human glioblastomas, we produced a driver-gene wide dataset of gene-associated, functional regulatory elements. Many genes were controlled by cis-regulatory networks composed of multiple regulatory elements, each of them providing a defined positive or negative input to the overall regulatory output of the network. Surprisingly, DNA methylation induces enhancers and silencers to acquire new activity setpoints within wide ranges of potential regulatory effects, varying between strong transcriptional enhancing to strong silencing. Extensive analysis of methylation-expression associations reveals the organization of domain-wide cis-regulatory networks, and highlighted key regulatory sites which provide pivotal contributions to the network outputs. Consideration of these effects through mathematical models of gene expression variations signified prime molecular events underlying cancer-genes misregulation in hitherto unexplained tumors. Of the observed gene-malfunctioning events, gene misregulation due to epigenetic retuning of networked enhancers and silencers dominated driver-genes mutagenesis, compared with other types of mutation including coding or regulatory sequence alterations. Elucidation of this gene-transformation mechanism may open the way for methodological disclosing of the driving forces behind cancers and other diseases.

## Background

Gene misregulation processes were hypothesized to underlie genetic ‘dark matter’ diseases lacking sufficient numbers of mutations in the coding sequences of their driving genes (Vogelstein et al. 2013). Searches for regulatory mutations were frequently focused on promoter and enhancer sequences (Sur et al. 2012; Hnisz et al. 2013; Mansour et al. 2014; Mifsud et al. 2015; Hnisz et al. 2016; Weischenfeldt et al. 2017; Bahr et al. 2018), but in many cases the cause of tumor development was not revealed (Consortium 2020; Rheinbay et al. 2020). We hypothesized that co-analysis of genetic and epigenetic variations at enhancers and silencers may help understanding the driving events which underlie these unexplained diseases. Transcriptional silencers are DNA sequences that upon binding of repressors or co-repressors reduce the transcription potential of linked promoters (Brand et al. 1985; Donda et al. 1996; Ogbourne and Antalis 1998; Burke and Baniahmad 2000; Zeng et al. 2021). However, regulatory loci may swap between enhancing and silencing effects on transcription in alternate cellular contexts (Grass et al. 2003; Rosenbauer et al. 2006; Huang et al. 2008; Gisselbrecht et al. 2020; Ngan et al. 2020). The molecular mechanisms which control these alternations are not fully understood. In line with the ability of regulatory sites to mediate enhancing or silencing effects, silencer loci were associated with both activating and repressing transcription factors and chromatin states, across cellular conditions (Kolovos et al. 2012; Qi et al. 2015; Huang et al. 2019; Doni Jayavelu et al. 2020; Pang and Snyder 2020; Cai et al. 2021; Huang et al. 2021). Silencers and enhancers were showed to cooperate in the regulation of gene transcription (Huang et al. 2019), but without thorough understanding of the mechanism and the factors that guide the mode of action of regulatory sites and the cooperation between them, it has been impossible to characterize the effect on normal and abnormal gene activities.

To deal with this challenge, we developed a method for detection and annotation the organization, activities and the interactions of silencers and enhancers in cancer tumors. DNA methylation is a sensitive and quantitative indicator of regulatory activity levels (Wiench et al. 2011; Aran and Hellman 2013), therefore a potentially effective marker for the mapping and elucidation of regulatory structures. However, the puzzling effects of methylation on both activation and silencing of gene expression have complicated the interpretation and utilization of methylation data. In particular, whereas methylation of gene promoters generally denotes gene repression (Bergman and Cedar 2013), both positive and negative effects on expression were observed among non-promoter regulatory sites (Gellersen and Kempf 1990; van Eijk et al. 2012; Wan et al. 2015). The mechanism underlying these opposing effects on transcription were not explained. Positive methylation-expression associations may reflect methylation-mediated silencing of repressor genes which in turn promote the expression of trans-controlled genes, or the hyper-methylation of transcribed gene bodies due to coupling between methylation and transcription apparati (Hellman and Chess 2007; Aran et al. 2011). Such secondary effects can be efficiently detected by analyzing inter-genic expression interactions. Alternatively, prime regulatory mechanisms may underlie positive methylation-expression associations, e.g., methylation-driven elimination of transcriptional repressors from silencers, or methylation-driven promotion of activator binding to enhancers. While evidence for direct effects of DNA methylation on transcriptional enhancers has been presented (Charlet et al. 2016; Wang et al. 2018), its effect on silencers and/or on the interaction between silencers and enhancers remains unknown. With so many levels of complication it is clear why, to date, DNA methylation information has not been utilized to its fullest capacity.

Here we demonstrate that integrative methylation and functional analysis may resolve this complexity, and by that allows efficient utilization of DNA methylation data to elucidate critical regulatory mechanisms. Building on this new platform, we describe an essential molecular mechanism which drives harmful gene activities in a common human disease.

## Results

### Producing integrative genetic-epigenetic maps of cis-regulatory domains

We developed a strategy for methylation-centered interrogations of functional gene-associated regulatory elements. While the method is applicable to many genes and diseases, this study focused on 125 pan-cancer and/or glioblastoma (GBM) driver genes, and 52 reference genes (Supplemental Table 1). To focus on regulatory sites that may alternate their mode of action across tumors, we initially evaluated the regulatory inputs provided by Histone 3 mono-methylated Lysine 4 (H3K4me1)-marked sites among various types of cancer. Clearly, H3K4me1 sites showed similar frequencies of positive and negative associations between methylation and expression levels (Supplemental Fig. 1A). Moreover, many of these sites switch between positive and negative effects on expression of the given genes, across cancers (Supplemental Fig. 1B). Based on this results, we targeted loci that carry H3K4me1 marks, and also the activity marker H3K27ac in some (but not all) of subjected glioblastoma tumors (Materials and Methods). An analysis of normal and cancer brains showed relative enrichment of DNase hypersensitivity signals within the targeted chromatin regions (Supplemental Table 2), thus confirmed their regulatory potential. Many of the target genes were not firmly assigned to particular topologically-associated domains (TADs) (Supplemental Fig. 2). Therefore, we choose to allocate all putative cis-acting regulatory elements within two million-base pair (Mbp) windows around the target gene promoters, thus ensuring unbiased evaluations of gene-associated sites within equivalent genomic spans. RNA probes (n=38,050, 120bp each) were designed for all CpG methylation sites (n=140,494) within these chromatin blocks (Supplemental Tables 3 and 4). By targeting the RNA probes to GBM tumors across patients with age, gender and GBM-subtype ranges characteristic of this disease (Supplemental Table 5), we obtained libraries of captured DNA segments (Supplemental Table 6 and Supplemental Fig. 3) representing the spectrum of sequence and methylation variations of the tumors. These libraries served as input material for parallel analyses of the regulatory function and the gene association status of the targeted loci (Fig. 1).

**Fig. 1.**
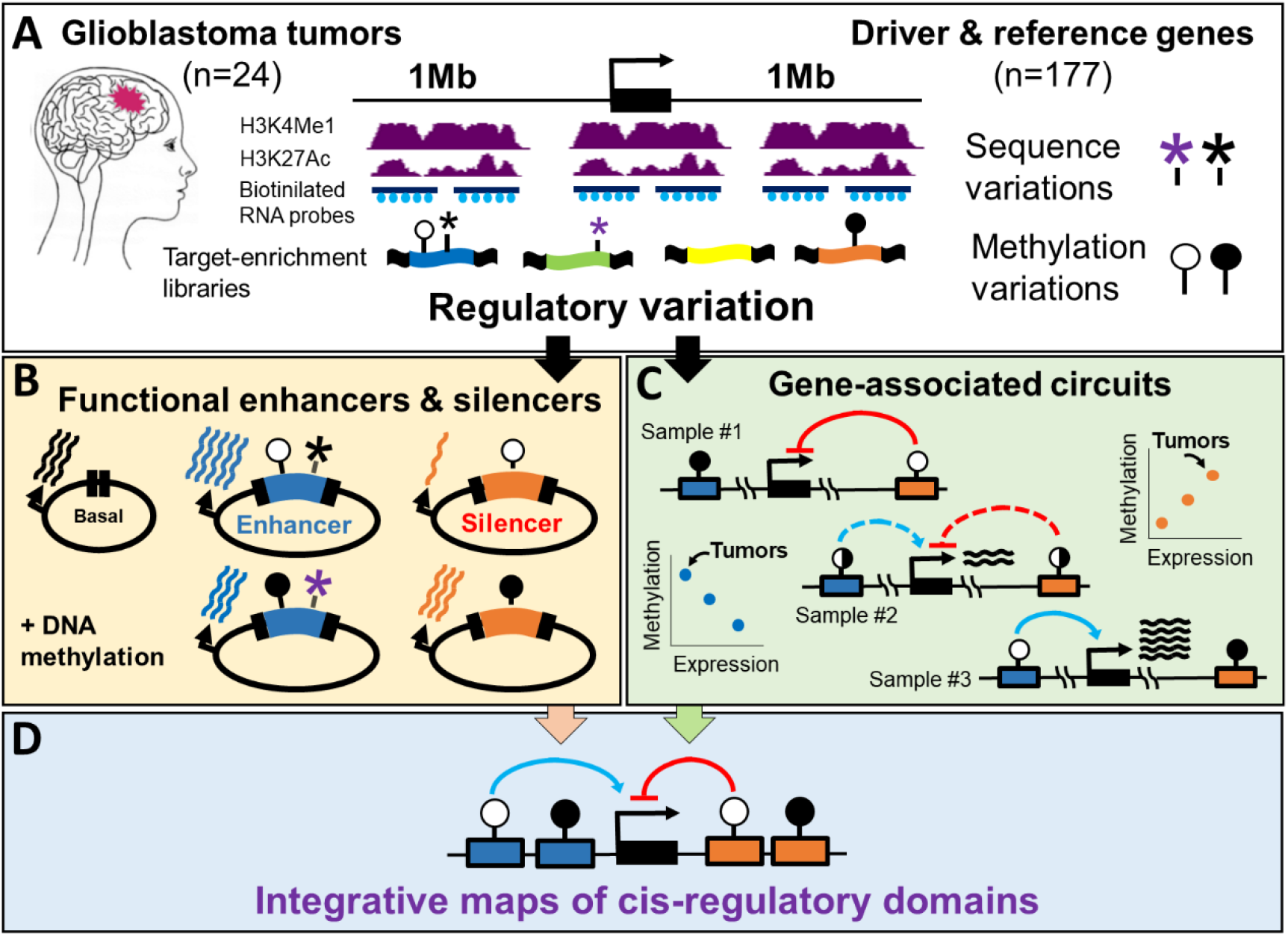
Methylation-centered interrogation of functional gene-associated regulatory elements. **(A)** Regulatory chromatin blocks were identified among glioblastoma (GBM) tumors in 2-Mb regions surrounding 125 driver and 52 reference cancer genes. H3K4Me1-marked/H3K27ac-variable chromatin segments encompassing methylation and sequence variations were captured from GBM tumor biopsies using biotinylated RNA probes. The obtained target-enriched libraries, representing the spectrum of GBM regulatory variation, were used for functional annotation of the targeted regions before or after DNA methylation **(B)**, or subjected to deep bisulfite sequencing providing methylation-site resolution of gene-associated positive and negative regulatory circuits **(C)**. The integration of functional and gene-associated data allows disclosing of cis-regulatory structures **(D)**.

### Silencers and enhancers are co-distributed along gene domains

Functionality of the captured regulatory elements was examined in GBM cells, using a massively paralleled reporter assay adapted for detection of silencers and enhancers (Materials and Methods). Transcriptional activity score (TAS) analysis revealed 26,152 significant (q<0.05) regulatory elements along the targeted gene domains, of them 9,204 silencers and 16,948 enhancers (Fig. 2A-C, Supplemental Fig. 4). The evaluation of additional 16,030 targeted genomic elements showed no significant functions. Analysis of the chromatin around the annotated elements in a variety of other cell types, showed that the loci annotated as silencers or as enhancers in GBM cells shared the characteristics of open, TF-bound regulatory chromatin (Fig. 2D). In most (176 of 177) of the analyzed gene domains we observed multiple (11-693) functional regulatory elements. Of these domains, 175 contained both enhancers and silencers (Supplemental Fig. 5). We concluded that regulatory elements are similarly distributed between enhancer and silencer functionalities across regulatory gene domains of GBM cells.

**Fig. 2.**
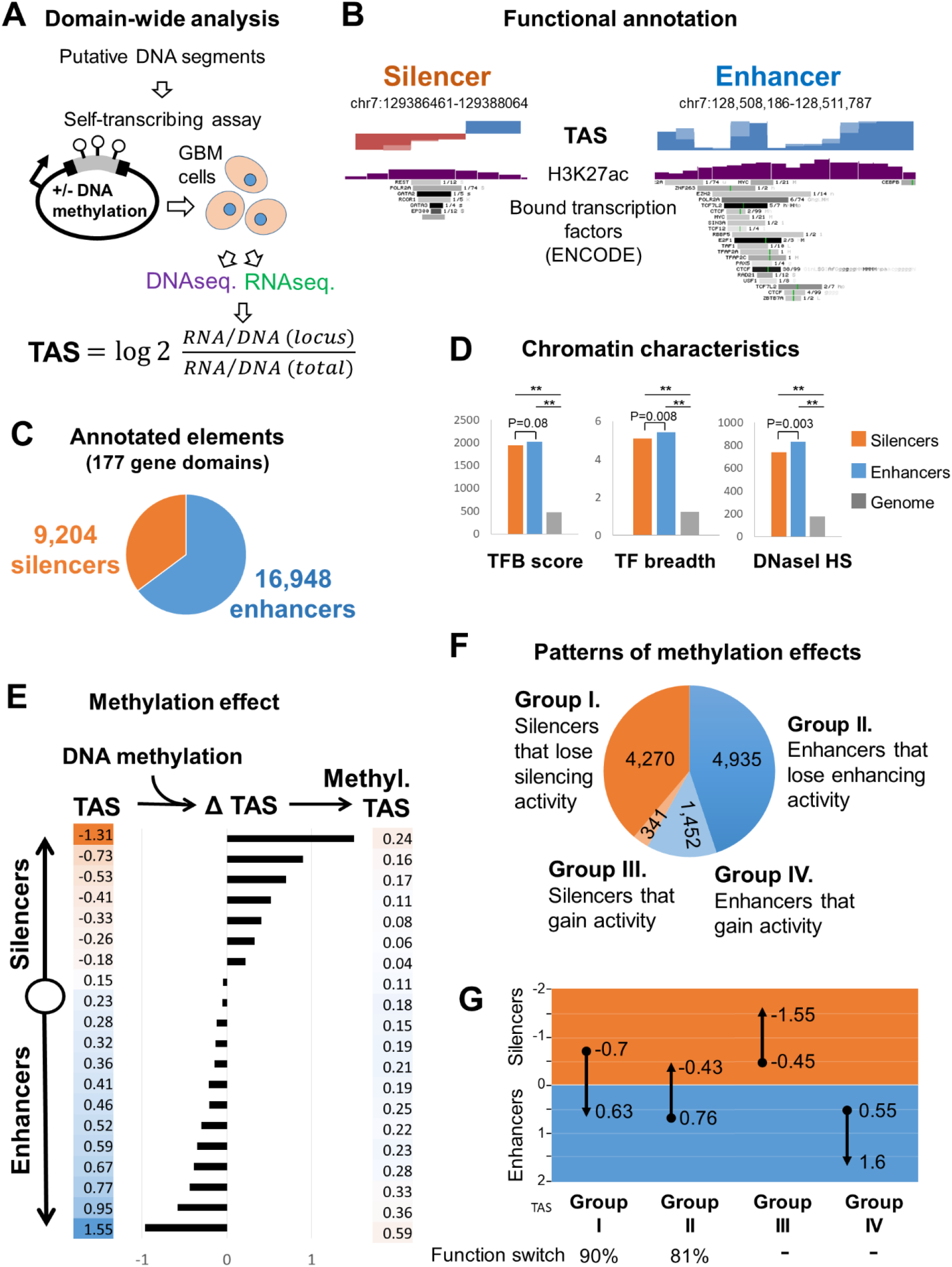
DNA methylation modify the transcriptional effect of enhancers and silencers. **(A)** Method: Putative regulatory DNA segments were captured from GBM tumors and allowed to drive self-transcription in T98G GBM cells, following complete de-methylation or after in-vitro methylation of the expression vector. Local DNA to RNA ratios, relative to the total DNA to RNA ratio, denotes transcriptional activity score (TAS) of the evaluated DNA segments. **(B)** Maps of example genomic regions containing enhancers and silencers. Local activity scores are shown as positive (blue) or negative (orange) bars. H3K27ac bars denote the fraction of the analyzed GBM tumors which displayed this marker of active regulatory chromatin. Bound TFs in a variety of different cell types are given as a reference for the general regulatory activity of the regions **(C)** Frequencies of regulatory elements that annotated as functional silencers or enhancers along the targeted gene domains. **(D**) Regulatory chromatin characteristics of enhancer and silencer loci. Level of transcription factors binding (TFB), factor variety (breadth), and DNase I hyper-sensitivity are shown. The data showed TF binding and chromatin accessibility across a variety of different cell types (ENCOD data). **(E)** Effect of DNA methylation on equal-number groups of regulatory elements ranked by TAS. **(F)** Patterns of methylation effects on silencers and enhancers. **(G)** Effect of DNA methylation on TAS level of the regulatory groups shown in panel **F**. The arrow heads indicate TAS level post-methylation. Fractions of sites that switched activities are given below. ** p<1E-20

### DNA methylation induces silencers and enhancers to acquire new activity set points

Instructive effects of methylation was next examined by comparing the transcriptional outputs of reporter genes derived by un-methylated versus methylated cis-regulatory elements (Supplemental Fig. 4). Of the 26,152 annotated regulatory elements, 10,998 displayed ≥1.5-fold TAS differences between methylated and un-methylated states. The other 15,154 (57.9%) elements may be insensitive to methylation, or affected below the detection threshold of the assay. Of the silencers and enhancers that displayed detectable effects, the majority (83.7%) reduced their original activities, or even gained the opposing functionality (i.e., enhancers became weaker or turned silencers, and vice versa), upon methylation (Fig. 2E-G). However, 16.3% of the methylation-responding elements showed the opposite effect, i.e. increased regulatory activity upon DNA methylation (Fig. 2F). Overall, methylation induces many enhancers and silencers to change their regulatory function. Many of them switched between enhancing and silencing modes (Fig. 2G), however, the effect of methylation was not restricted to complete switching between full enhancing and full silencing functionalities. Rather, it allowed silencers and enhancers to adopt new activity set points within ranges of enhancing to silencing effects, possibly by affecting the balance between bound activators and repressors. In line with this possibility, the analyzed regulatory elements bind a repertoire of activators and repressors across cell types, regardless of their functional annotation in GBM (Supplemental Fig. 6). We concluded that core regulatory sequences may retuned on their operative scales between enhancing and silencing inputs to the transcriptional machinery. DNA methylation is apparently required and sufficient to induce these effects in GBM cells.

### Methylation data reveals the organization of cis-regulatory units and networks

Utilizing the same capturing libraries that were used for the functional assays, we next studied methylation-expression associations in intact GBM genomes. We analyzed the correlation between the methylation levels of the captured sites and expression levels of the targeted genes among 24 GBM samples (Fig. 3A, Supplemental Table 1), as described (Aran and Hellman 2013; Aran et al. 2013). The resulted correlation maps revealed associations between certain regulatory sites and controlled genes, termed the cis-regulatory circuits of the genes. After filtering out 197 gene-body and promoter sites to exclude possible secondary effects (Supplemental Fig. 7), 900 regulatory circuits of 109 genes were obtained. Of them, 42% denoting positive relationships with expression, and 58% negative (Supplemental Table 7). Most (78%) of the genes had multiple (2-68) circuits, averaging 8.3 (3.5 positive, 4.8 negative) circuits per gene. Clusters of gene-associated methylation sites formed defined regulatory units of tens to thousands (average 834, median 333) bp-long spans containing up to 31 homogenous (positive or negative) sites. Each of these units mediate positive or negative input to the transcription of a particular gene (Supplemental Table 8). Merging of association and functional data revealed cis-regulatory structures, in which positive units aligned with silencers, and negative units with enhancers (Fig. 3B).

**Fig. 3.**
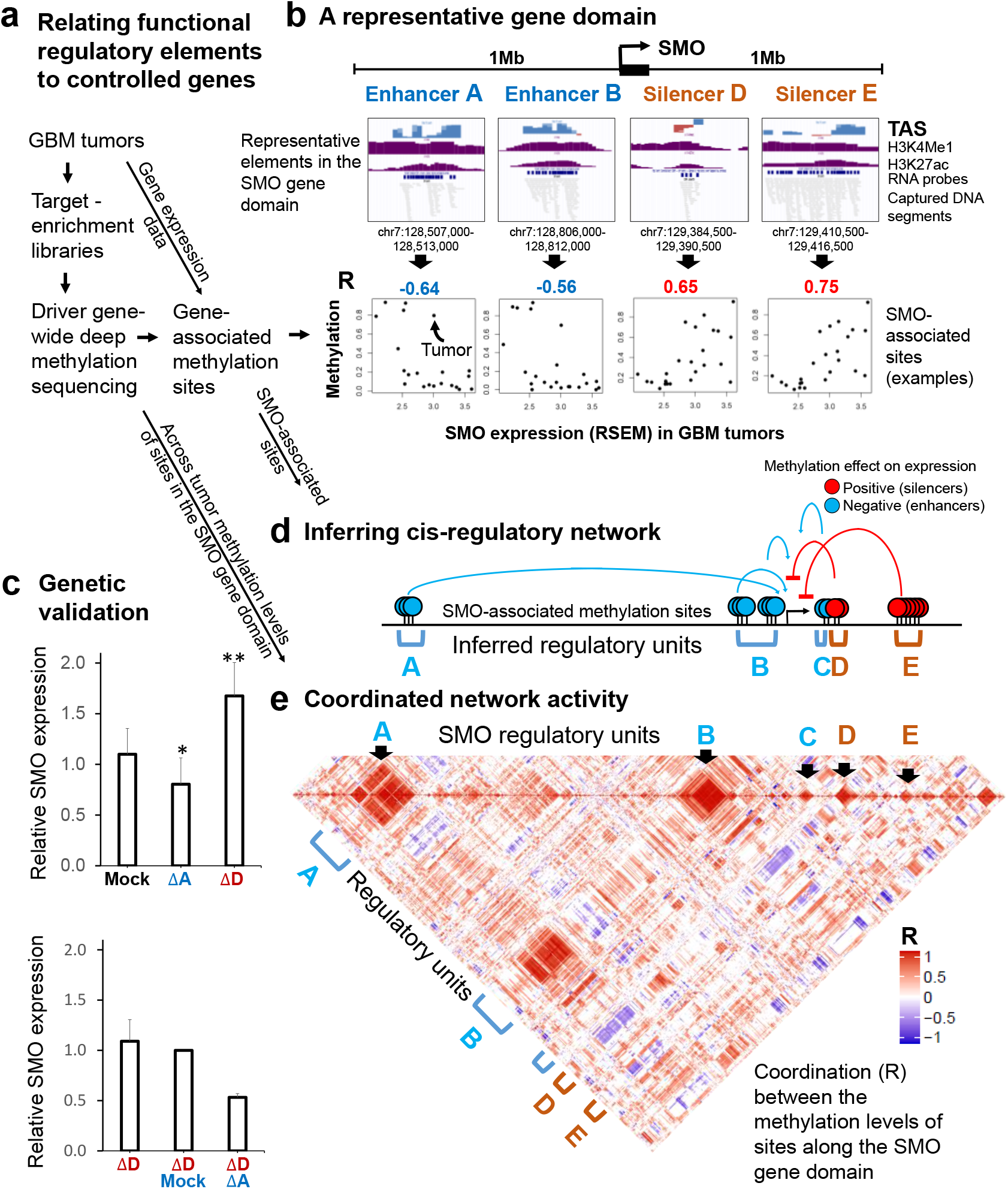
Methylation-based deciphering of cis-regulatory networks in bona fide tumor chromatin. **(A)** Methylation-based association of regulatory sites with controlled genes. **(B) Top:** Examples of functional enhancer and silencer elements that identified along the SMO driver gene domain through the massively parallel assay presented in figure 2. Even-sized windows (about 20X larger than the median size of regulatory units) are shown. **Bottom:** Correlations between DNA methylation and SMO expression levels across GBM tumors, for representative methylation sites in the functional elements. **(C)** Validation of the predicted effects of SMO regulatory units via manipulations of GBM genomes. **Top:** Effects of enhancer or silencer deletions versus mock genomic targeting by scrambled targeting guides. Units are labeled as in panel A. **Bottom:** Effect of enhancer deletion on the background of a silencer deletion. Bars represent standard deviations based on >= four biological replications. **(D)** Gene-associated sites reveal networks of homogenous, positive or negative regulatory units that cooperatively control SMO expression variation. **(E)** A diagram showing the correlation between the methylation level of each methylation sites in the SMO domain, and the methylation levels of all other sites in the domain, across 24 GBM tumors. In the tumors with the highest expression of the gene, enhancers were unmethylated and silencer were methylated, and vice versa. * p<0.05; ** p<0.005

### Genomic editing experiments verify silencing and enhancing inputs

We applied genomic manipulation experiments to verify particular predictions of the functional gene-association annotations. The Smoothened, Frizzled Class Receptor (SMO) driver-gene, as example, was abnormally expressed in 23 of the 24 tumors. Three functional enhancers and two functional silencers, consisting 29 associated methylation sites, structure the cis-regulatory network of the gene (Supplemental Table 7). Indeed, removing of a functional, SMO-associated enhancer from the genome of GBM cells reduced SMO expression relative to mock-treated cells, whereas deletion of a silencer unit gained its expression. Moreover, deletion of the enhancer unit has similar effect on the wild-type and silencer-deletion backgrounds (30-50% reduction relative to the background expression levels), suggesting that the enhancer and the silencer units provide additive inputs to the transcriptional machinery (Fig. 3C, Supplemental Fig. 8 and 9).

### Co-residing genes controlled by independent cis-regulatory networks

Interestingly, networked silencers and enhancers were reversely coordinated across the tumors, so tumors with lower expression of a given gene tend to display unmethylated silencers and methylated enhancers, whereas tumors with higher expression of the gene displayed the opposite arrangements (Fig. 3D-E, Supplemental Fig. 10, Supplemental Data 1). Moreover, networks of different genes were independent of each other, even when their units were spatially intermixed (Supplemental Fig. 11, Supplemental Data 2), suggesting that neighboring genes are controlled by gene-specific cis-regulatory networks spanning overlapped genomic domains.

### Mathematical modulation signifies key regulatory sites

We further explored the interaction between networked silencers and enhancers by examining multiplexed effects on gene expression: Given a certain effect of an arbitrarily selected regulatory site on expression of its controlled gene, we asked whether multiplexed models that also consider other associated sites provide improved expression prediction. Therefore, redundant regulatory sites should provide no improvement, whereas antagonists or synergistic sites are expected to improve the prediction provided by each of the sites alone. Using stepwise analyses, we identified the best models of possible combinations of up to four sites (Fig. 4A). For example, the eighteen TNFAIP3-associated sites produced predictive R-values ranging between −0.72 and 0.71 for each individual site (Supplemental Table 7). The tests of the 4,029 possible combinations of one to four sites out of the 18 revealed a model that incorporated the methylation levels of two positive and two negative sites, providing better prediction power than each of the sites alone (R=0.9, p=1.41E-06). Hence, the revealed model signifies the methylation sites providing the best description of the gene expression-variation. By that, it hints to the particular regulatory sites, out of all associated sites, which are most significant to the regulation of the gene. Similarly, the best model for the SMO gene, incorporating the methylation level of two positive and two negative sites, provided better prediction power (R=0.8, p=0.00027) than each of the 29 associated sites alone. As in the case of TNFAIP3, these sites resided within positive and negative regulatory units (Fig 4B). Note that the model used no preliminary assumptions regarding the nature of the most predictive sites. Therefore, the fact that both positive and negative sites were called, suggests that they are jointly response to the determination of gene expression level.

**Fig. 4.**
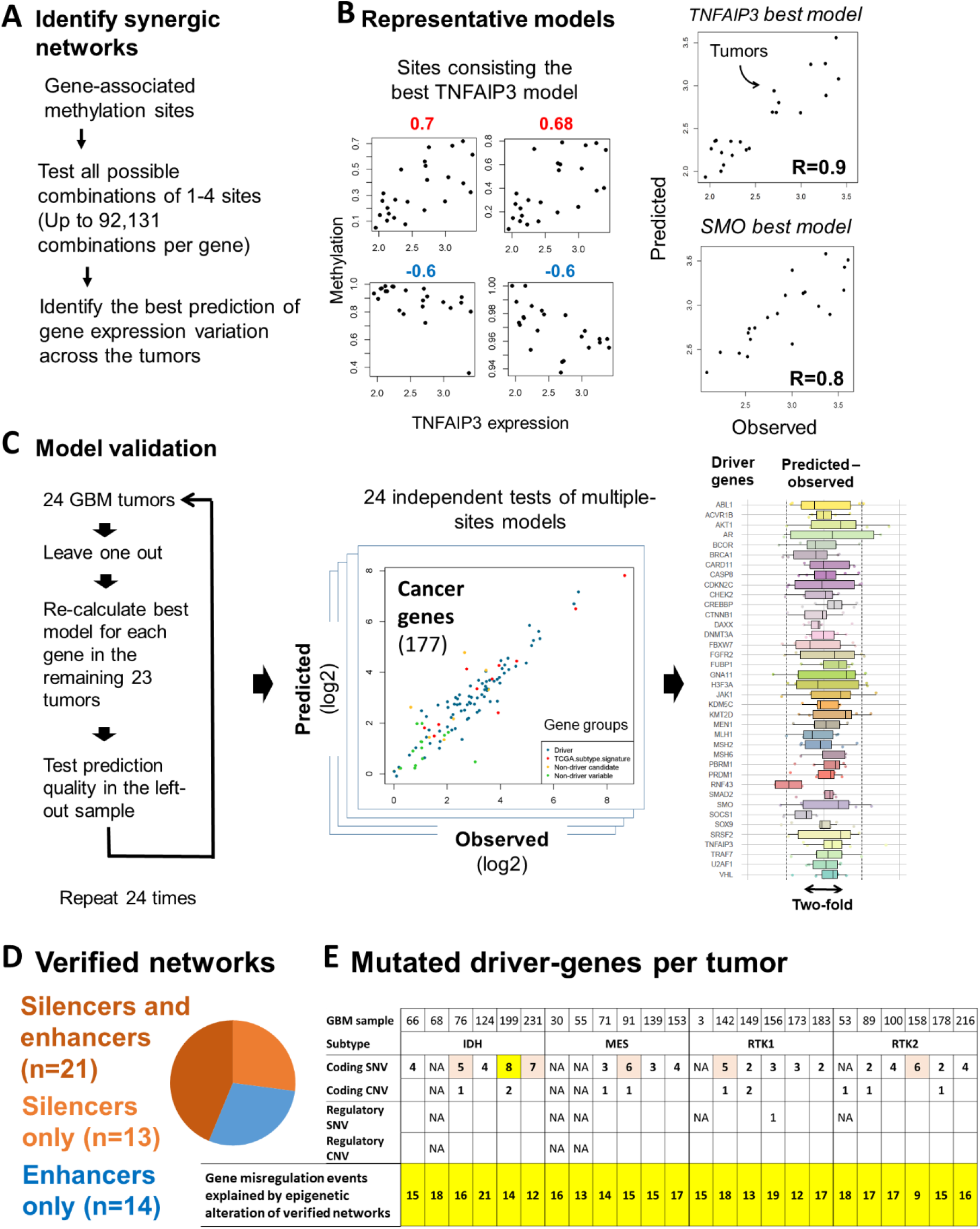
Networks of epigenetically-tuned transcriptional silencers and enhancers govern disease driver-gene malfunction. **(A)** Development of methylation-based models of gene expression variation. **(B)** Example models. **Left:** methylation versus expression of the sites consisting the best prediction model of the TNFAIP3 gene. **Right:** predicted versus observed variation of the TNFAIP3 and the SMO genes across the tumors. SMO model was based on the four sites shown in figure 3b. **(C)** Verification of the mathematical models for the driver genes. Verifications on non-driver models are given in Supplemental Fig. 12. **(D)** Participation of silencers and enhancers in confirmed Cis-regulatory networks. **(E)** Numbers of driver-gene per tumors that affected by sequence or methylation mutations in their coding or regulatory components. **SNV:** Single Nucleotide Variation; **CNV:** Copy Number Variation. Misregulated genes are genes that display >2 fold expression deviation from normal brain in the given tumor sample. Highlighted cell indicates tumors with at least five (orange) or eight (yellow) mutated driver-genes.

Overall, out of 105 genes with significant models, the expression of 58 genes were best predicted by synergic combinations of sites, providing better prediction than each of the sites alone (Supplemental Table 9). The power of mathematically-significant models were further verified by testing their predictions in tumors that were not used during the model development (Fig. 4C, Supplemental Fig. 12, Supplemental Data 3). Of the 48 genes with validated synergic or validated single-site models, silencers were involved in the regulation of 34 genes (Fig. 4D).

We concluded that mathematical modulation of methylation effects provides a way to identify contributing regulatory sites and to explore their roles in synergic, gene-specific networks. The analysis efficiently uncovered important cis-regulatory sites and networks, out of the many gene-associated sites and numerus possible combinations of sites presented in gene regulatory domains.

### Epigenetically-retuned cis-regulatory networks guides gene transformation

Finally, we compared the contribution of silencers, enhancers and coding mutations to driver gene malfunction. In the majority (68.4%) of the tumors, fewer than five driver genes were affected by nonsynonymous or copy number mutations (Fig. 4E), in line with previous analyses of this cancer (Kandoth et al. 2013; Ceccarelli et al. 2016). To reveal the effect of regulatory sequence mutations we deep-sequenced the uncovered silencers and enhancers in eight of the patients, and analyze the effect of sequence variations on expression of the associated genes. Notably, only one possible event revealed, aside from common sequence polymorphisms (Supplemental Note 1). As current models of cancer predict a minimum number of five to eight mutated driver genes (Vogelstein et al. 2013), regulatory and coding sequence mutations alone cannot explain the appearance of a majority of the GBM tumors. In contrast, all tumors included more than eight abnormally expressed driver genes that associated with methylation-tuned regulatory units and explained by confirmed methylation-based models of expression variations (Fig. 4E). Silencers were involved, alone or in cooperation with enhancers, in almost two-thirds of these misregulation events, and were implicated in the malfunction of genes driving a wide range of cancer initiation and progression processes (Table 1, Supplemental Fig. 13). We conclude that epigenetically-retuning of networked regulatory elements plays a prime role in the malfunction of cancer driver-genes.

**Table 1.**
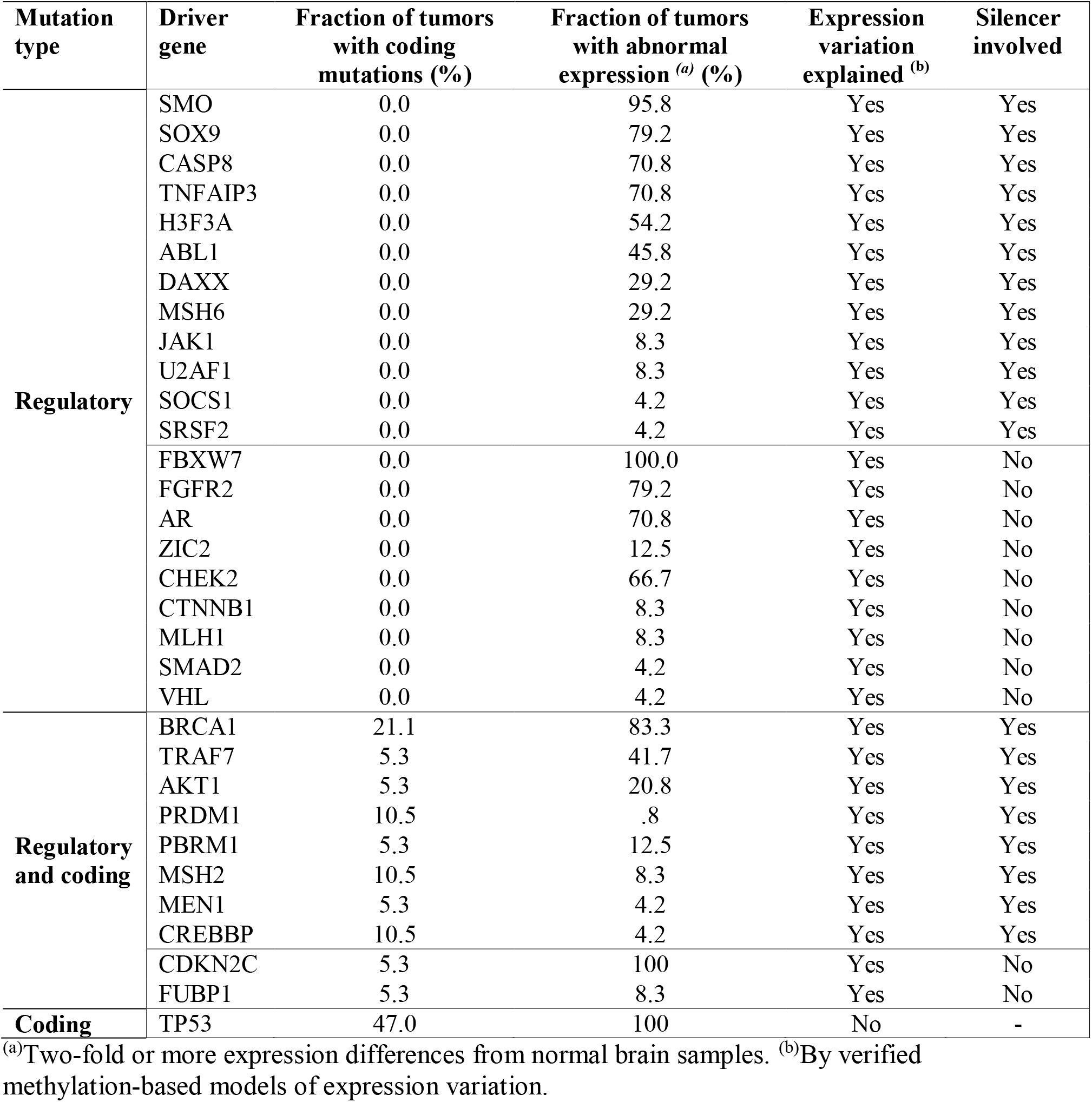
Genes affected by regulatory or coding mutation.

## Discussion

Limited knowledge of cis-regulatory mechanisms hampered discrimination and understanding of causative regulatory mutations in human diseases. Here we revealed a pivotal role for methylation-guided re-setting of cis-regulatory outputs in malfunctioning of cancer-driving genes.

In lines with former reports (Grass et al. 2003; Rosenbauer et al. 2006; Huang et al. 2008; Doni Jayavelu et al. 2020; Gisselbrecht et al. 2020), many of the regulatory elements that were mapped in this study were able to switch their functions between enhancing and silencing of transcription. The observation that silencer and enhancer loci may bind both activating and repressing transcription factors (Supplemental Fig. 6), suggests that this enhancer-silencer functional switching may mediated by alternation in the balance between bound activators and repressors. We further showed that functional resetting of regulatory elements is not limited to binary enhancing-or-silencing modes. Rather, ranges of activity set points may be acquired. Under experimental conditions, DNA methylation appeared required and sufficient to retune the level and the mode of regulatory activities (Fig. 2). In bona fide cancer tumors, knowing the level of DNA methylation at key sites of cis-regulatory networks was sufficient to obtain very effective predictions of gene expression variations. The alternations of DNA methylation levels at these sites explain the malfunctions disease driver genes, which were not associated with sequence alternations (Fig. 4). Taken together, these experimental and in-vivo evidence support a prime role for methylation-mediated resetting of cis-regulatory networks in cancer gene malfunction. Through consideration of these effects, one may uncover key mutational events underlying cancers and other diseases.

A key feature of our experimental design is the co-detection of enhancing and silencing effects. Though early massively paralleled reporter assays used expression vectors which are not sensitive to the detection of silencers (Muerdter et al. 2018), it was recently shown that adapted vectors may overcome this bias (Doni Jayavelu et al. 2020). In line with this report, we showed that effective detection of enhancing and silencing effects may achieved by applying vectors with a certain level of basal transcription.

Our study focused on particular chromatin regions within the regulatory domains of cancer driver-genes. Since in published glioblastoma data (Johnston et al. 2019), HiC-mapped TADs were not defined for about a third of our genes of interest, we choose to target regulatory sites up to 1 million bp from the gene promoters. This provided full coverage of >80% of the mapped TADs while allowing analysis of all targeted genes. Within these domains, we focused on methylation sites with H3K4me1 and variable H3K27ac-marked chromatin. We showed that these targeting criteria are relevant to a large bulk of glioblastoma silencers and enhancers. Nevertheless, additional enhancers and silencers of the targeted genes may reside outside of the areas covered in this study.

Assembling of the produced methylation-expression data in mathematical models of gene-expression variation revealed key determinants of gene regulation. We showed that gene-specific networks, spanning large genomic regions, are composed of relatively small regulatory units (median size=333bp, Supplemental Table 8) of defined silencing or enhancing activities. We also found that co-residing genes may be independently controlled by spatially-overlapped gene-specific, cis-regulatory networks. The spatial organization of genes with their regulatory elements, and the way by which neighboring genes may be independently regulated within shared regulatory domains, were been poorly understood. Our data suggest that gene-specific regulatory patterns may achieved even if the gene and its regulatory sites were not isolated within gene-specific territories (Sun et al. 2019). Instead, we showed that multiple gene-specific networks may use shared genomic spans (Supplemental Fig. 11, Supplemental Data 2). Wiring the regulatory units of these overlapping networks with their targeted gene promoters is topologically complex, unless if occurs in separated time windows. While direct imaging of such temporal arrangements is still unavailable, our discovery of coordinated network-specific activities between distant regulatory units in human tumor DNA (Fig. 3E, Supplemental Data 1), and similar findings in the mouse (Li et al. 2019), support this view.

Finally, we found that methylation-modified networks of silencers and enhancers provide a prime component of GBM gene malfunction (Table 1). The integration of silencer and enhancer methylation mutations complete the missing effect of coding and regulatory sequence mutations (Consortium 2020; Rheinbay et al. 2020) and explains the appearance of tumors with small-then-expected number of altered driver-genes. The high prediction power provided by the identified silencer and enhancer sites flags them as candidate biomarkers for RNA-free tumor profiling. Whether genetic or epigenetic editing of these sites will improve GBM phenotypes, and the relevance of our findings to other illnesses, remains to be determined.

## Methods

### GBM samples and data

Tumor biopsies and associated clinical data were collected and encoded at the DKFZ Institute, Heidelberg, Germany. Whole-genome, whole-exome, H3K4me1 and H3K27ac chromatin immunoprecipitation (GSE121719) and RNA sequencing of the GBM biopsies and the normal brain samples (GSE121720), and the analyses of coding DNA mutation, gene expression and DNA copy number variation, were performed at the DKFZ. Encoded de-personalized DNA samples and data were used as input materials for target enrichment of gene regulatory regions and associated DNA methylation and non-coding DNA mutation analyses, which were performed at the Hebrew University, Jerusalem, Israel.

### Genes

Genes analyzed in the study included the pan-cancer driver genes listed by Vogelstein et al.(Vogelstein et al. 2013) including the GBM driver genes listed by Kandoth et al.(Kandoth et al. 2013), but excluding the HIST1, H3B and CRLF2 genes due to missing expression data, and the AMER1 gene for which probe design failed. Cancer type-specific genes (n=23) were selected from a published list of 840 genes (Verhaak et al. 2010). Non-driver candidate GBM genes (n=14) suggested by BR. Non-driver variable genes (n=22) were defined as those showing top expression variation among the 70 analyzed GBM samples for which we found at least two correlative sites in the TCGA-GBM dataset. We used the genomic coordinates for gene features from the hg19 refGene table of the UCSC Genome Browser (Karolchik et al. 2014)

### Public databases

#### The Cancer Genome Atlas (TCGA)

Gene expression (RNAseqV2 normalized RSEM) and DNA methylation data (HumanMethylation450) were download in May 2019 using TCGAbiolinks(Colaprico et al. 2016; Silva et al. 2016; Mounir et al. 2019) for the following cancer types: BRCA (778 genomes), CESC, (304), COAD (306), ESCA (161), GBM (50), KICH (65), KIRC (320), KIRP (273), LIHC (371), LUAD (463), PAAD (177), SKCM (103), THYM (119).

#### NIH Roadmap Epigenomic Project (Roadmap Epigenomics et al. 2015)

H3K4me1 broad peaks of corresponded TCGA tumor types and DNaseI cell specific narrow peaks of normal brain (E081 and E082).

#### Encyclopedia of DNA Elements (ENCODE)(Consortium 2012)

DNaseI hyper-sensitivity peak clusters (wgEncodeRegDnaseClusteredV3.bed.gz) and transcription factor ChIP-seq clustres (wgEncodeRegTfbsClusteredWithCellsV3.bed.gz) and DNase brain tumors data (Gliobla and SK-N-SH). The ENCODE transcription factor binding (TFB) scores presented in Fig. 2 represent the peaks of transcription factor occupancy from uniform processing of ENCODE ChIP-seq data by the ENCODE Analysis Working Group. Scores were assigned to peaks by multiplying the input signal values by a normalization factor calculated as the ratio of the maximum score value (1000) to the signal value at one standard deviation from the mean, with values exceeding 1000 capped at 1000. Peaks for 161 transcription factors in 91 cell types are combined here into clusters to produce a summary display showing occupancy regions for each factor and motif sites within the regions when identified. One-letter code for the different cell lines is given in https://hgsv.washington.edu/cgi-bin/hgTrackUi?hgsid-2654998_o9Di2gB797ixpn7O898j4DsMV3Ro&g=wgEncodeRegTfbsClusteredV3.

#### Additional public data

HiC Data for TADs were downloaded from https://wangftp.wustl.edu/hubs/johnston_gallo/ (Johnston et al. 2019)

### Cell lines

Human GBM T98G cells were purchased from the ATCC collection (ATCC^®^ CRL-1690™), and cultured in minimum essential medium-Eagle (Biological Industries), supplemented with 10% heat-inactivated FBS #04-127-1A (Biological Industries), 1% penicillin/streptomycin P/S # 03-031-1B (Biological Industries), 1% L-glutamine #03-020-1C (Biological Industries;), 1% non-essential amino acids, #01-340-1B (Biological Industries) and 1% sodium pyruvate #03-042-1B (Biological Industries), at 37°C and 5% CO_2_.

### Target enrichment assays

Variable regulatory regions were defined as the regions carrying H3K4me1 marks in all tumors, and also H3K27ac in at least 25% of the tumors, but not in at least other 25% of the tumors. RNA probes were designed to target all methylation sites within these regions, utilizing the SureDesign tool (https://earray.chem.agilent.com/suredesign/). Probe duplication was applied in cases (n=8,652) of >5 CpG sites within the 120 bp span of the probes. Repetitive regions were identified by BLAT(Bhagwat et al. 2012) and excluded from the design. Custom-designed biotinylated RNA probes were ordered from Agilent Technologies (https://www.agilent.com). Enrichment libraries of GBM-targeted regulatory DNA segments were constructed using the SureSelect protocol #G9611A (Agilent) for Illumina multiplexed sequencing, which used 200 nanograms genomic DNA per reaction, or the SureSelect Methyl-Seq protocol #G9651A using one microgram genomic DNA per reaction. Quality and size distribution of the captured genomic segments were verified by TapStation nucleic acids system (Agilent) assessments of regular or bisulfite-converted libraries. Target enrichment efficiency and coverage was evaluated via sequencing.

### Massively paralleled reporter assay

Massively parallel functional assays were performed as described by Arnold et al (Arnold et al. 2013), with the following modifications:

#### Reporter backbone

The pGL3-promoter vector (Promega, GenBank accession number U47298) was modified as described in Supplemental Fig. 14.

#### Genomic inputs

Plasmid libraries were constructed using a target-enriched library as input material: One microliter of adaptor-ligated DNA fragments from the AK100 target enrichment library was amplified in eight independent PCR reactions, using KAPA Hifi Hot Start Ready Mix #KK2601 (KAPA Biosystems). Reaction conditions included 45 seconds (s) at 95°C, 10 cycles of 15s at 98°C, 30s at 65°C, 30s at 72°C, and 2 min final extension at 72°C, applying the forward Ilumina universal primer: 5′-TAGAGCATGCACCGGTAATGATACGGCGACCACCGAGATCT-3′ and reverse Indexed Ilumina primer: 5′-GGCCGAATTCGTCGACCAAGCAGAAGACGGCATACGAGAT-3′, containing Illumina adapter sequences. A specific 15nt extension was added to each adapter as homology arms for directional cloning. PCR reactions were pooled and purified on NucleoSpin Gel and PCR Clean-up #740609 columns (Macherey-Nagel). The screening vector was linearized with AgeI-HF and SalI-HF restriction enzymes (NEB) and purified through electrophoresis and gel extraction. Purified PCR products were cloned into the linearized vector by recombination with the adapter-ligated homology arms in 12 reactions of 10 μl each, applying the In-Fusion HD #639649 kit (Clontech). The reactions were then pooled and purified with 1x Agencourt AMPureXP DNA beads #A63881 (Beckman Coulter) and eluted in 24 μl nuclease-free water.

#### Library propagation

Aliquots (n=12, 20 μl each) of MegaX DH10B T1 Electrocomp Bacteria #C640003 (Invitrogen) were transformed with 2 μl of the plasmid DNA library, according to the manufacturer’s protocol, except for the electroporation step, which was performed using the Nucleofactor 2b platform (Lonza) Bacteria program 2. Every three transformation reactions were pooled (total of 4 reactions) for a one-hour recovery at 37°C, in SOC medium, while shaking at 225 rpm, after which, each reaction was transferred to 500 ml LB AMP (Luria Broth Ampicillin) for overnight 37°C incubation, while shaking at 225 rpm. Propagated plasmid libraries were extracted using the NucleoBond Xtra Maxi Plus Kit (#740416) (Macherey-Nagel). To verify unbiased amplification of the targeted genomic segments, size distribution and coverage of the library were analyzed before and after the propagation step (Supplemental Fig. 15).

#### *In-vitro* methylation assay

Complete de-methylation stages achieved by propagation of the libraries in bacteria following PCR amplification stages. *In-vitro* methylation of the de-methylated plasmid DNA was performed using the New England Biolabs CpG Methyltransferase M.SssI #M0226M according to the manufacturer’s instructions. Efficient methylation level was confirmed by using a DNA protection assay against FastDigest HpaII #FD0514 (Thermo Scientific) digestion (Supplemental Fig. 16)

#### Transfection to GBM cells

20 μg of DNA were transfected into 2×10^6^ T98G and U87 cells at 70-80% confluence, using the Lipofectamine 3000 transfection kit #L3000-015 (Invitrogen), according to the manufacturer’s protocol. In each experiment, 5×10^7^ cells were transfected and incubated at 37°C, for 24 h.

Isolation of plasmid DNA and RNA from GBM cells: Plasmid DNA was extracted from 2.5×10^7^ cells, 24 h post-transfection. Cells were rinsed twice with PBS pH 7.4, using the NucleoSpin Plasmid EasyPure kit #740727250 (Macherey-Nagel), according to the manufacturer’s protocol. Total RNA was extracted from 2.5×10^7^ cells 24h post-transfection using GENEZOL reagent # GZR200 (Geneaid), according to the manufacturer’s protocol. The polyA+ RNA fraction was isolated using Dynabeads Oligo-(dT)25 #61002 (Thermo scientific), scaling up the manufacturer’s protocol 5-fold per tube, and treated with 10 U turboDNase #AM2238 (Invitrogen) at 20 ng/μl 37°C, for 1 h. Two reactions of 50 μl each, were pooled and subjected to RNeasy MinElute clean up kit #74204 reaction (Qiagen) to inactivate turbo DNase and concentrate the polyA+ RNA.

#### Reverse transcription

First strand cDNA synthesis was performed with 1-1.5 μg polyA+ RNA in a total of four reactions, 20 μl each, using the Verso cDNA Synthesis Kit #AB1453B (Thermo scientific) according to the manufacturer’s protocol, with a reporter-RNA specific primer (5′-CAAACTCATCAATGTATCTTATCATG-3′). cDNA (50 ng) was first amplified by PCR, at 98°C for 3 min, followed by 15 cycles at 95°C for 20s each, 65°C for 15s, 72°C for 30s. Final extension was performed at 72°C for 2 min, using the Hifi Hot Start Ready Mix (KAPA), with reporter-specific primers. Forward primer: 5′-GGGCCAGCTGTTGGGGTG*T*C*C*A*C-3′ which spans the splice junction of the synthetic intron, and reverse primer: 5′-CTTATCATGTCTGCTCGA*A*G*C-3′, where “*” indicates phosphorothioate bonds. In total, 16-20 reactions were performed. The amplified products were purified with 0.8 x Agencourt AMPureXP DNA beads and eluted in 20 μl nuclease-free water. The resultant purified products served as a template for a second PCR performed under the following conditions: 95°C for 3 min, 12 cycles of 98°C for 15s, 65°C for 30s, 72°C for 30s. Final extension was performed at 72°C for 2 min, with forward Ilumina universal primer: 5′-TAGAGCATGCACCGGTAATGATACGGCGACCACCGAGATCT-3′ and reverse Indexed Ilumina primer: 5′-GGCCGAATTCGTCGACCAAGCAGAAGACGGCATACGAGAT-3′. PCR products were purified with 0.8 x Agencourt AMPureXP DNA beads, eluted in 10 μl nuclease-free water, and pooled.

### Transcriptional activity analysis

Quality and size distribution of extracted plasmid DNAs and RNAs were verified using TapeStation (Supplemental Fig. 17). DNA and cDNA samples were sequenced using the HiSeq2500 device (Illumina), as per the 125bp paired-end protocol. Alignment with the hg19 reference genome was performed on the first 40 bp from both sides of the DNA segments, using Bowtie2 (Langmead and Salzberg 2012). Reads with mapping quality value above 40 aligned with the probe targets were considered for further analyses. Each of the captured genomic segments was given a unique ID according to genomic location and indicated the total number of DNA and RNA reads. Only on-target segments with at least one RNA read (n= 623,223 pre-methylation; 304,998 post-methylation) were included. >99% of the targeted regions were presented following the propagation in bacteria and re-extraction from T98 cells (Supplemental Table 10). Technical and biological replications performed using illumina MiSeq sequencing.

Transcriptional activity score (TAS) was calculated as follows:

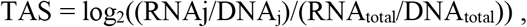

where j is a genomic element and RNA_total_ or DNA_total_ is the sum of all segment reads.

For the analyses of isolated regulatory elements, TAS was determined in 500bp, 50% overlapping windows, across the genome, based on the DNA and RNA reads of segments overlapping with the given window. TAS significance was tested by Chi-square against total RNA to DNA. Multiple comparisons were corrected. Functional regulatory elements were defined as elements with FDR q value < 0.05 and minimum 100 RNA reads, where positive TASs were defined as enhancers, and negative as silencers. The methylation effect was analyzed by calculating TAS difference between treatments, where regulatory elements with a difference of ≥1.5-fold activity were counted.

### Inferring cis-regulatory circuits

#### Methylation sequencing

Methyl-seq-captured libraries were sequenced using a Hiseq2500 device (Illumina), by applying paired-end 125bp reads. Sequence alignment and DNA methylation calling were performed using Bismark V0.15.0 software (Krueger and Andrews 2011) against the hg19 reference genome. The sequencing yielded 52-149 million reads per sample, at an average mapping efficiency of 78.1%, average bisulfite efficiency of 97.6%, and 99.4% on target average. Overall, a mean coverage of 916 reads per site was obtained, and 86% of the targeted sites were covered by at least 100 reads. Sites that reported in less than eight of the tumors were excluded from the analyses.

#### Circuit annotation

Correlation between the expression level of each targeted gene and the DNA methylation level of targeted CpG sites in a 2Mbp region flanking its transcription start site (TSS), was assessed by applying pairwise Spearman’s rank correlation coefficient with Benjamini-Hochberg correction for multiple-hypothesis testing at FDR < 5%. Sites that correlated (R^2^ > 0.1) with expression of the PTPRC (CD45) pan-blood cells marker, were considered a possible result of blood contamination and were eliminated from later analyses, as described (Aran et al. 2016). Potential secondary effects were considered in two cases (1) The correlated site was included within the prescribed portion (the gene body, excluding the first 5Kbp) of another gene; (2) The correlated site was located within the promoter (from TSS-1500bp to TSS+2500bp) of another gene. For these cases, a correlation between the expression level of the genes was tested, and circuits with R^2^>0.1 that fit one of the scenarios described in Supplemental Fig. 7, were excluded.

#### Methylation-based prediction of gene expression

For each gene, we tested all the possible combinations of one to four associated sites. For each combination with full data in at least 12 tumors, we generated a predictive model of expression level based on multiple linear regression of the sites methylation levels. A significant model (q value < 0.05), evaluated by ANOVA for Linear Model Fit, and corrected for the number of possible models per-gene by FDR. A gene was considered to have a synergic model if the predictive value of the model was better than each of the involved sites alone.

Validation of methylation-based predictions was performed using the leave-one-out cross validation approach for assessing the generalization to an independent data set. One round of cross-validation involves 23 data sets (called training set) in which performing all the analysis, and one sample for validating the analysis (called testing set). The cross-validation was performed x24 times. For each training data set, cis-regulatory circuits were generated (as described in Circuit annotation sub-section above) and possible predictive models were developed for the targeted genes. Prediction quality of each gene was then tested in the 24 rounds, by comparing predicted versus observed expression level. Difference up to 2 fold considered as success. The ability to accurately predict the expression level of a gene was considered verified if has good prediction quality in at least 20 of the 24 rounds.

### Analysis of coding sequence variations

VCF files describing single nucleotide variations (SNV) were provided by the DKFZ. Synonymous SNV, SNVs overlapping with published SNPs (COMMON), or SNVs with a less than 25-read coverage or bcftools-QUAL score >20, were excluded. Copy number variations (CNV) were analyzed by whole-genome sequencing (WGS) data provided by the DKFZ. Association between gene expression and copy number was evaluated by Pearson or Spearman’s correlations. p-values were adjusted for multiple-hypothesis testing using the Benjamini-Hochberg method, with FDR < 5%.

### Analysis of regulatory sequence variations

#### Pre-alignment processing

GBM tumors (n=8) were sequenced using the paired-end 250-or 300bp read protocol in Illumina MiSeq V2 or V3 devices. FASTQ files were filtered, and sequence edges of Phred score quality >20 were trimmed up to 13 bp of Illumina adapter applying Trim Galore (http://www.bioinformatics.babraham.ac.uk/projects/trim_galore/). Reads that were shortened to 20 bp or less were discarded, along with their paired read. Exclusion of both reads was implemented after verifying that retention of unpaired reads did not significantly increase high quality alignment coverage. Quality control of the original and filtered FASTQ files was performed with FastQC (http://www.bioinformatics.babraham.ac.uk/projects/fastqc), deployed to verify the reduction in adapter content and the increase in base quality following the filtering stage. Duplicates were removed at the pre-alignment stage with FastUniq (Xu et al. 2012). Duplicate pair-ends were removed by comparing sequences rather than post-aligned coordinates, allowing preservation of variant information.

#### Sequence alignment

Sequences were aligned to GRCh37/hg19 assembly of the human genome applying paired-reads Bowtie 2(Langmead and Salzberg 2012). Discordant pairs or constructed fragments larger than 1000bp were discarded, thereby improving mapping quality by allowing both reads to support mapping decisions. Default values (Bowtie 2 sensitive mode) were applied to end-to-end algorithm parameters, seed parameters, and bonus and penalty figures. Outputted SAM and BAM alignment files were examined using the Picard CollectInsertSizeMetrics utility to verify correctness of final insert-size distribution (http://broadinstitute.github.io/picard. Version 1.119).

#### Variation calling

A BCF pileup file was generated from each BAM file using the samtools(Li et al. 2009) mpileup function, set to consider bases of minimal Phred quality of 30 and minimal mapping quality of 30. Variant calling performed using bcftools, was initially set to output SNPs only to create SNP VCF files, according to the recommended setting for cancer (Narasimhan et al. 2016). The VCF files were filtered by applying depth of coverage (DP) above 40 and statistical Quality (QUAL) above 10. DP filtering in this context refers to DP/INFO in the VCF file, which is a raw count of bases.

#### Variant post-processing

Post-processing of VCF SNPs included additional filtering, variant frequency calculation, mapping variants to probes and mapping variants to public databases, performed with a custom-written Python script. Additional depth coverage filtering of 20 was applied on the high-quality bases, which were selected by bcftools as appropriate for allelic counts. Frequency calculations were based on high-quality allelic depth (ratio of each allelic depth to sum of all allelic depths). SNPs were mapped to the following dbSNP(Sherry et al. 2001) and ClinVar(Landrum et al. 2018) databases: dbSNP/common version 20170710, dbSNP/All version 20170710 and clinvar_20170905.vcf. A match was determined when the position, reference and variant were all in agreement. In our analysis, we refer to de-novo variations (not in COMMON and not in ALL) which were detected in at least one sample (of eight). For each targeted gene, we counted the number of de-novo variations that were at a distance of ±500bp from its correlated sites.

#### Regulatory CNVs

Non-coding CNVs were detected from WGS of 5Kbp sliding blocks in a 2Mbp region flanking gene TSSs, with a 50% overlap. Correlation of the total copy number (TCN) of each block with the gene expression level was assessed (at least six samples with available TCN data, Pearson and Spearman correlation). Correlation p values were adjusted for multiple-hypothesis testing using the Benjamini-Hochberg method..

### Genome editing

#### Design and cloning of sgRNA

Guides to perturb SMO regulatory units were designed using the ChopChop, E-CRISP and CRISPOR softwares. 20-bp sgRNA sequences followed by the PAM ‘NGG’ for each unit, were identified and synthesized. For the SMO regulatory unit at chr7:128,507,000-128,513,000 designated unit “A”, 4 guides were cloned into a backbone vector bearing Puromycin resistance (addgene, 51133), using the Golden Gate assembly kit (NEB^®^ Golden Gate Assembly Kit #E1601). Each guide sequence was cloned with its own U6 promoter and was followed by a sgRNA scaffold. For the regulatory unit at chr7:129,384,500-129,389,500, designated unit “D”, two guides were cloned into the same backbone plasmid using the same method (Supplemental Fig. 8).

#### Transfection/CRISPR-Cas9-mediated deletion

After validating the sgRNA sequences by Sanger sequencing, T98G or T98GdeltaSMO-D cells were co-transfected with a Cas9-bearing plasmid (addgene, 48138) and either the plasmid bearing the guides targeting SMO A, the plasmid with bearing the guides targeting SMO D, or the same plasmid harboring a nontargeting gRNA sequence (scramble), as a negative control. The molar ratio between the transfected guide plasmid and the Cas9 plasmid was 1:3, in favor of the plasmid not carrying the antibiotic resistance. 1.5-3*10^5 cells/ml, >90% viable, were plated one day prior to transfection in a 6-well dish. On the transfection day, each well received 3 microliter Lipofectamine^®^ 3000 Reagent, 5 microgram total plasmid DNA and 10ul of Lipofectamine^®^ 3000 Reagent (2:1 ratio). Puromycin (3 micrograms/microliter) was added to the cells one day after transfection. After 72h, the antibiotic was washed and the cells were left to expand. The cells were harvested 8-21d post-transfection and genomic DNA and RNA were immediately collected (Qiagen; DNeasy #69504 and RNeasy #74106, respectively).

#### Genotyping of mutant populations

Genomic DNA was subjected to genotyping PCR (primers listed in table). Deletion or partial deletion was confirmed by gel electrophoresis or TapeStation. RNA extracted from populations of cells bearing such mutations were then checked for an effect on SMO transcription level, using qPCR (QuantStudio 3 cycler, Applied Biosystems, Thermo Fisher Scientific, Waltham, MA, USA).

#### Single-cell dilution to obtain CRISPR-targeted cell clones

Puromycin-selected cells were isolated by trypsinization, counted and diluted to a concentration of 20 cells/100 microliters. Diluted cells (200 microliters) were then serially diluted, to ensure single-cell occupancy of rows 6-8 (eight dilution series). By calibrating the number of cells in the first row we ensured that single cells could be isolated from the sixth to eighth rows onwards. Cells were incubated until the low-density wells were confluent enough to be transferred to 24-, 12- and finally to 6-well plates. Selected clones were tested for a stable DNA profile and for SMO transcription level by genotyping PCR (primers listed in table), followed by gel electrophoresis or TapeStation and qPCR analysis, respectively.

#### RT-qPCR

Each isolated mRNA (500 ng) was transcribed to cDNA using the Verso cDNA Synthesis Kit (#AB-1453/A, Thermo Fisher Scientific, Waltham, MA, USA) according to provided instructions, using the oligo dT primer. qPCR was performed using the Fast SYBR™ Green Master Mix (#AB-4385612, Thermo Fisher Scientific, Waltham, MA, USA) and qPCR primers for SMO and reference genes HPRT and TBP (see table), on a QuantStudio 3 cycler (Applied Biosystems, Thermo Fisher Scientific, Waltham, MA, USA). The reaction was conducted in triplicates, and 20ng of template were placed in each well. For each primer set, a no-template control (NTC) was also run, to check for possible contamination. QuantStudio Design & Analysis Sofware v1.4.3 (Applied Biosystems, Thermo Fisher Scientific, Waltham, MA, USA) was used for analysis. All presented data were based on three or more biological replications of the genome editing experiments, each with three technical repeats of the DNA and RNA.

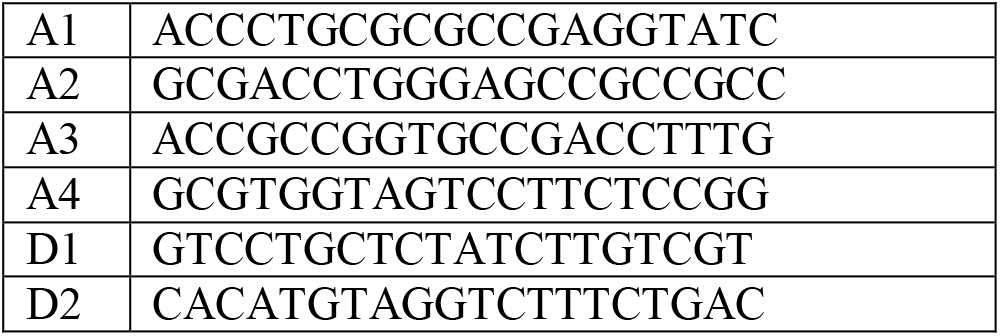
Guides list

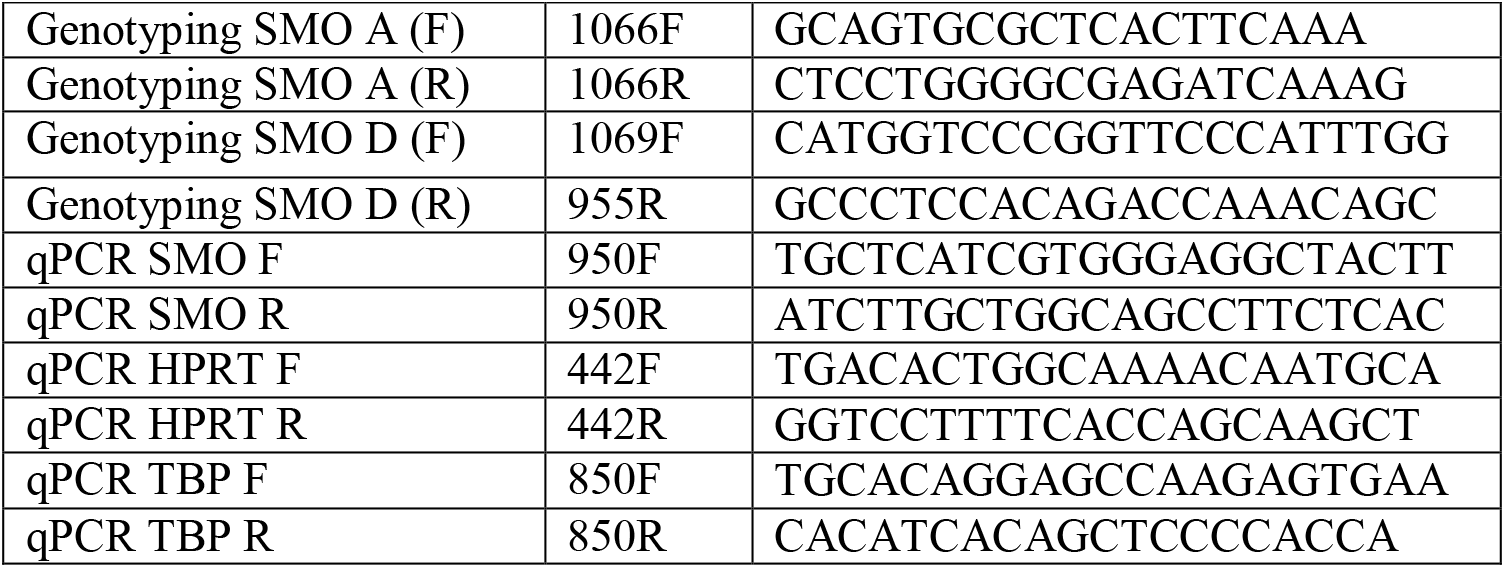
Primers list

## Data access

All raw and processed sequencing data generated in this study have been submitted to the NCBI Gene Expression Omnibus (GEO; https://www.ncbi.nlm.nih.gov/geo/) under accession number GSE163021 [secure token will be given upon request]. All other relevant data supporting the key findings of this study are available within the article and its Supplemental Information files.

## Acknowledgments

The European Research Council (ERC) Grant No. 724803 to A. H., the Israel Science Foundation Grant No. 948/16 to A.H., and a grant from the Heidelberg Center for Personalized Oncology (DKFZ-HIPO) to B.R. supported this work.

## Author contributions

Conceptualization, A.H.; Methodology, A.H. and Y.E.; Software, Y.E.; Formal Analysis, Y.E.; Investigation, R.L. and A.M.; Resources, B. R.; Data Curation, Y.E.; Writing - Original Draft, Y.E. and A.H.; Writing - Review & Editing, A.H. and B.R.; Visualization, A.H. and Y.E.; Supervision, A.H.; Project Administration, R.L., B.R., and A.H.; Funding Acquisition, A.H. and B.R.

