## Supplemental Material for "Methylation-mediated retuning on the enhancer-to-silencer activity scale of networked regulatory elements guides driver-gene misregulation"

Yifat Edrei, Revital levy, Anat Marom, Bernhard Radlwimmer and Asaf Hellman

**Supplemental Note 1**

**Supplemental Figure 1-17**

**Supplemental References 1-3**

**Supplemental Note 1**

In agreement with former glioblastoma analyses (Sturm et al. 2012; Mackay et al. 2017), analyses of copy number variations in the studied tumors showed moderate karyotype aberration (average tumor ploidy = 2.5) (Supplemental Table 5). Since copy-number alterations are not necessarily accompanied by abnormal expression of residing genes, we assessed functional events by examining the correlation between gene copy number and mRNA levels. Genes displaying significant correlations ($R^2$>0.3; $p<0.05$) between their copy numbers and expression levels across the tumors, and corresponding two-fold or greater expression deviation from normal brain tissue, were considered misregulated genes attributable to gene copy number variation (CNV). EGFR was the only gene that fully met these criteria. Possible subtle effects were observed for PDGFRA/KIT, MET and CHEK2. Overall, 0-2 genes per tumor were potentially altered by coding CNVs. To evaluate regulatory sequence mutations, single-nucleotide variations (SNV) were analyzed in the regulatory segments captured from eight of the GBM tumors. Only one potentially contributing variation reveled in 100-bp windows around gene-associated sites, and extended analyses using larger window sizes revealed no additional events. Theoretically, amplification or deletion of enhancer sequences may also affect transcription. However, a full screen of CNVs in 5,000 bp windows across the driver gene domains revealed no events apart from gene CNVs.

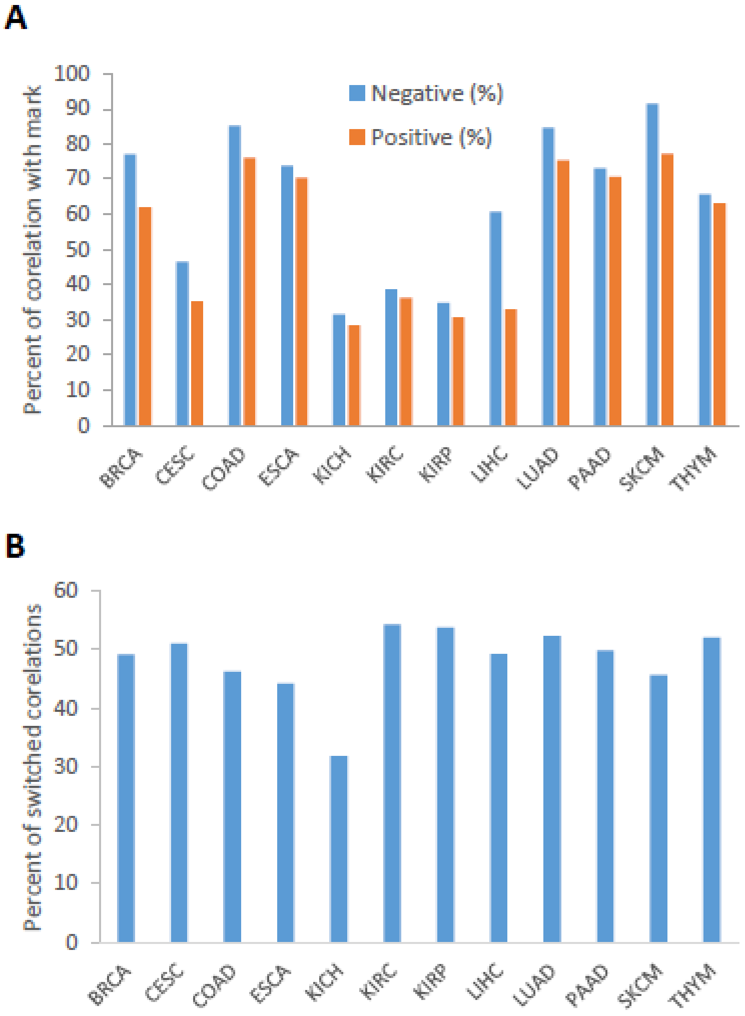


**Supplemental Figure 1. Methylation-expression associations in various cancer types. (A)** Percentages of negatively and positively-associated sites that carry H3K4me1 marks, out of all gene-associated sites across various types of cancer. The analysis performed on public TCGA data. **(B)** Percentages of gene-associated methylation sites in given types of cancers, which displayed the opposite effects on expression of the associated genes in at least one other cancer type.

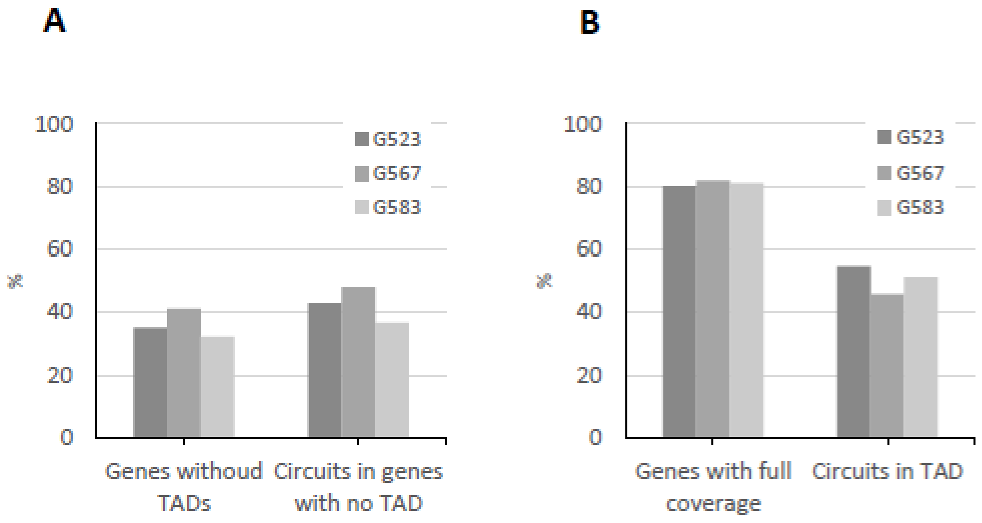


**Supplemental Figure 2. Overlapping between targeted gene domains (+/- 1Mb of TSS) and Hi-C-based topological associated domains (TAD). (A) Left:** Fractions of genes without TADs following Hi-C analysis of three GBM samples (25kb resolution) (Johnston et al. 2019). **Right:** Fractions of gene-associated sites that related to genes without TADs, out of all uncovered gene-associated sites. **(B) Left:** Fractions of genes with Hi-C-based TADs, for which our targeting criteria provide full coverage of the gene TAD. **Right:** Fractions of gene-associated sites within Hi-C-based TADs, out of all uncovered gene-associated sites.

**Sample #AK100 target enrichment library**

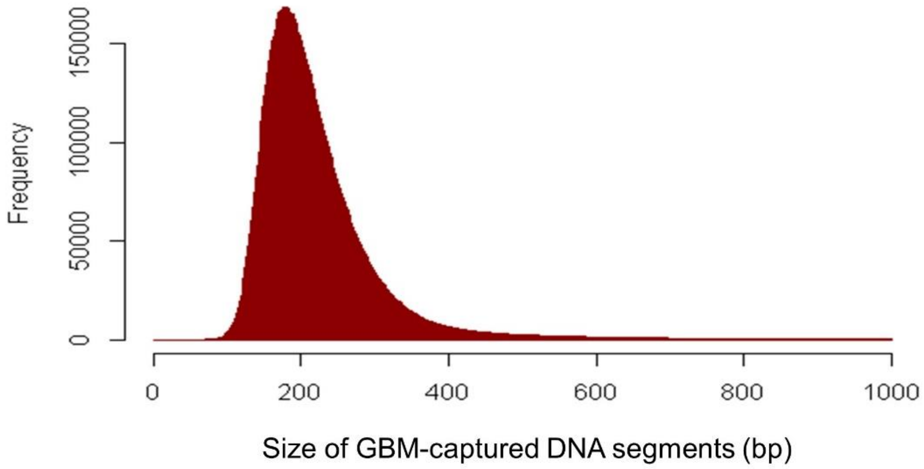


**Supplemental Figure 3.** A representative distribution of the size of DNA segments that were captured from the GBM tumors.

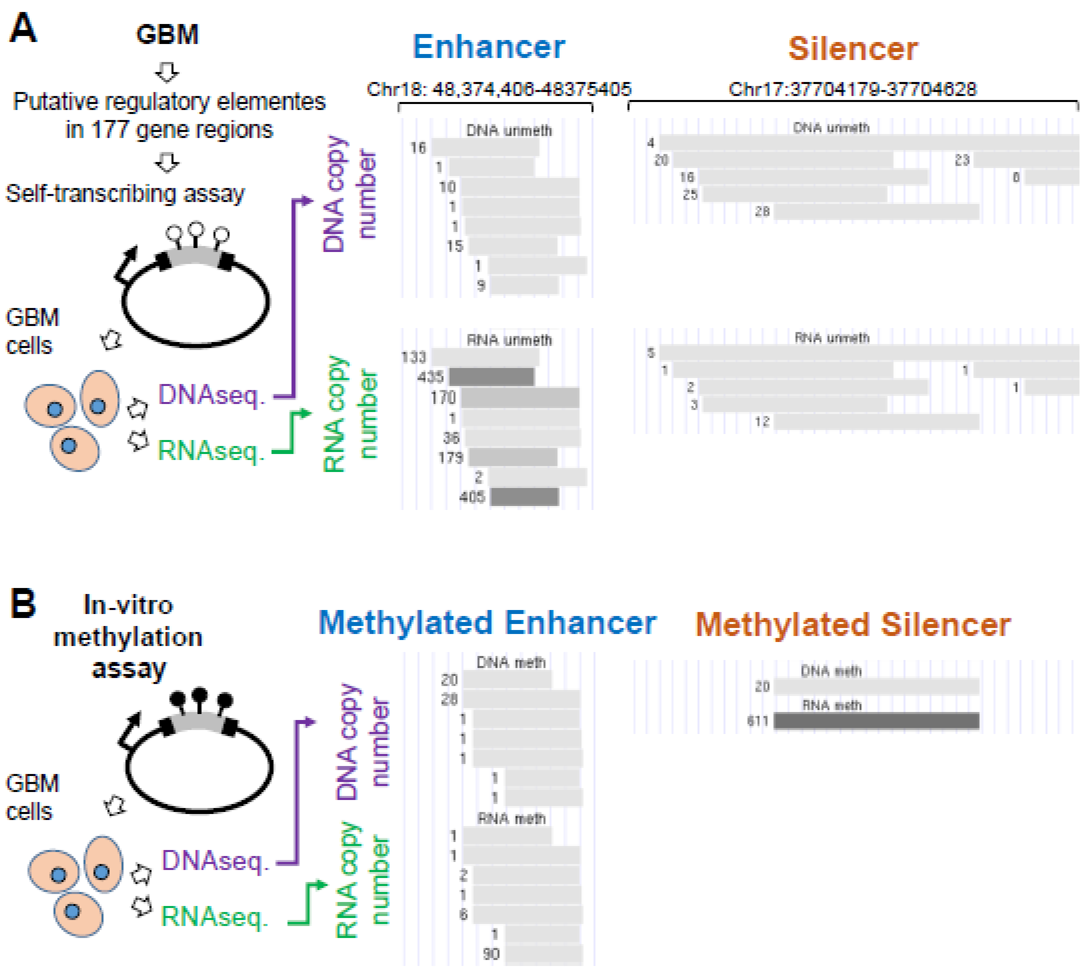


**Supplemental Figure 4. Functional annotation of isolated regulatory elements. (A)** Genomic segments (mean size=224bp) were captured from a GBM tumor, ligated downstream to minimal promoters and allowed to drive own transcription in T98G glioblastoma (GBM) cells. Plasmid DNA and RNA were then extracted from the GBM cells and sequenced. The ratio between DNA and RNA copy numbers, normalized to total DNA and RNA levels, denotes the transcriptional activity of the targeted elements. Example enhancer and silencer elements. DNA and RNA copy numbers are indicated to the left of each segment. **(B)** The enhancer and silencer shown in panel A following in-vitro DNA methylation.

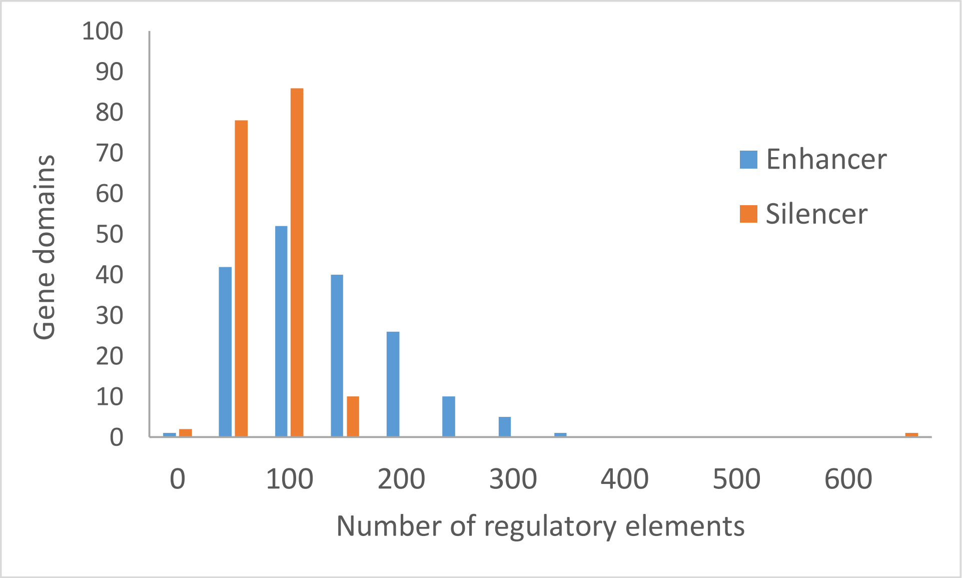


**Supplemental Figure 5**. Distribution of silencer and enhancer elements in the targeted gene domains.

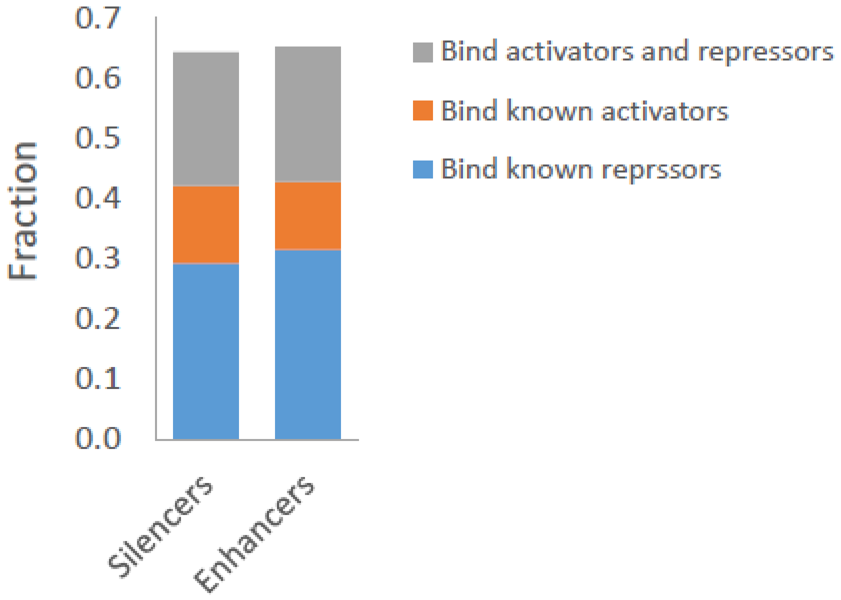


**Supplemental Figure 6**. Fractions of enhancers and silencers that bind activating, repressing, or both activating and repressing transcription factors across ENCODE cell lines. The list of activators includes: RNAP, GATA2, GATA3, EP300, BCL3, NFATC1, HNF4A, HNF4G, ELK4, ELK1 and IRF1. The repressors list includes: REST, YY1, ZBTB33, SUZ12, EZH2, RCOR1, CTCF, SMC3, RAD21, PAX5 and RUNX3.

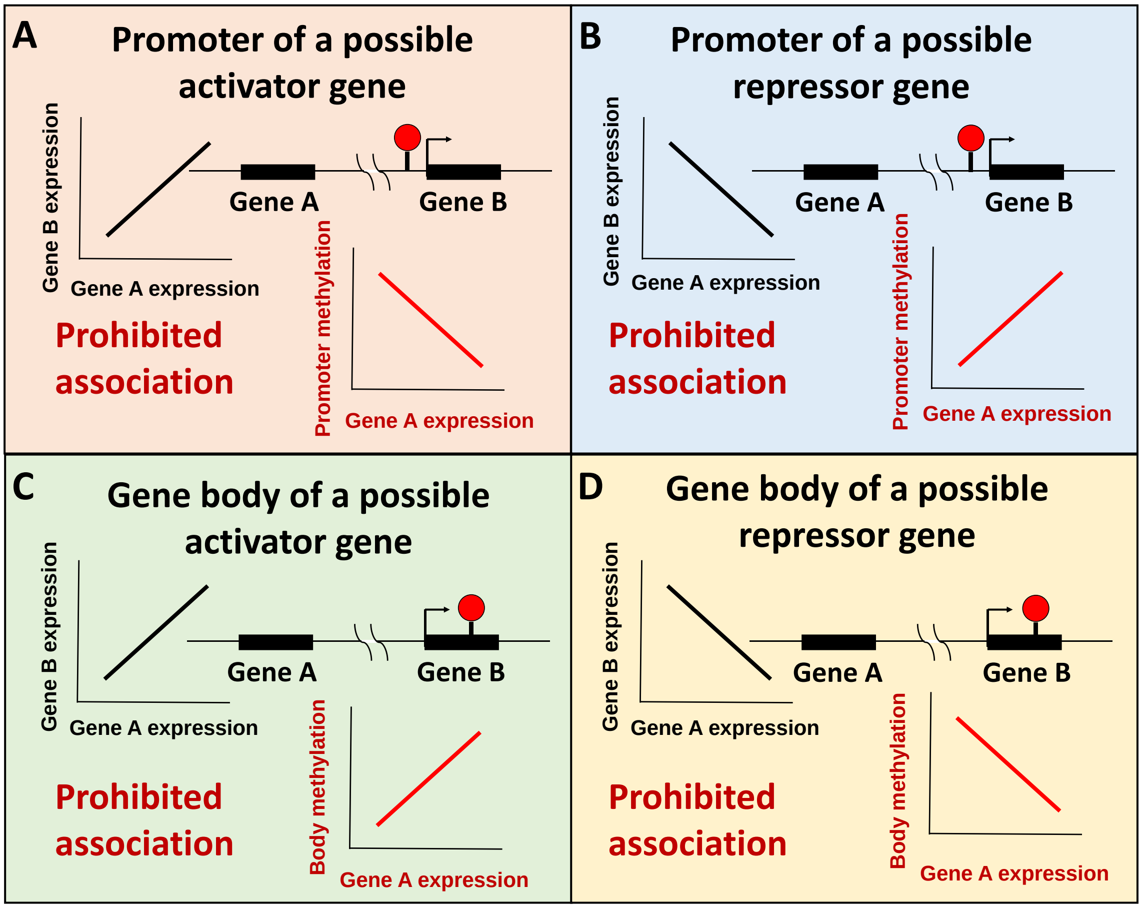


**Supplemental Figure 7. Eliminated associations due to possible secondary effects. (A)** Prohibited association between methylation of a promoter site and expression of a possible activator of the indicated gene A. **(B)** Prohibited association between methylation of a promoter site and expression of a possible repressor of the indicated gene A. **(C)** Prohibited association between methylation of a gene-body site and expression of a possible activator of the indicated gene A. **(D)** Prohibited association between methylation of a gene-body site and expression of a possible repressor of the indicated gene A

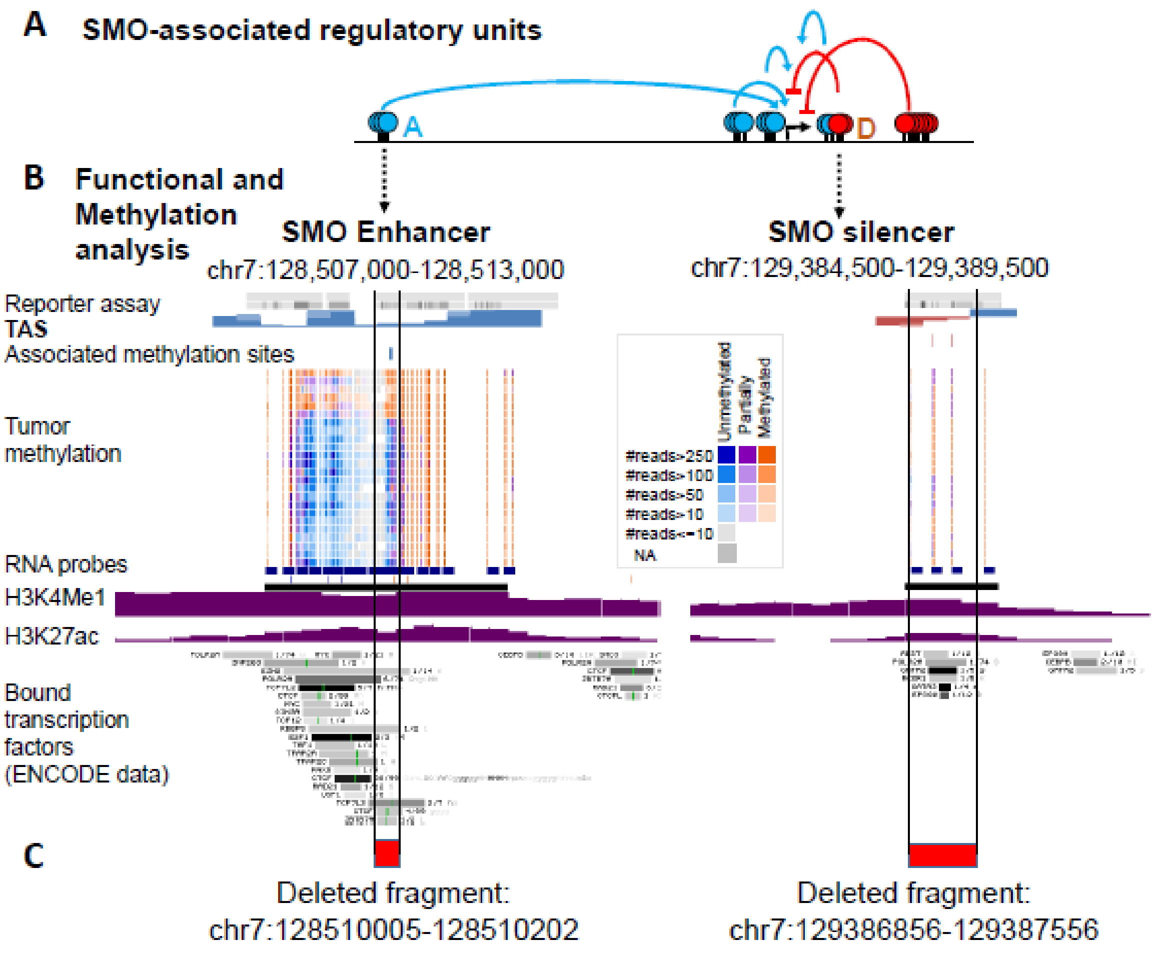


**Supplemental Figure 8**. **Alignment of positive and negative units with silencers and enhancers. (A)** A schematic map showing the five regulatory units of the SMO driver gene. **Blue:** negative methylation-expression associations. **Red:** positive associations. **(B)** Functional and methylation analyses of SMO enhancer and silencer units. **Top:** Transcriptional Activity Score (TAS) analyzed through reporter assay analysis. Blue**:** positive (enhancing) scores. Brown: negative (silencing) scores. **Middle:** DNA methylation levels of the 24 analyzed GBM tumors. **Bottom:** Chromatin marks and bound transcription factors. **(C)** Genomic coordination of the knockout regions in the genomic editing experiments (Fig. 3C and Supplemental Fig. 9).

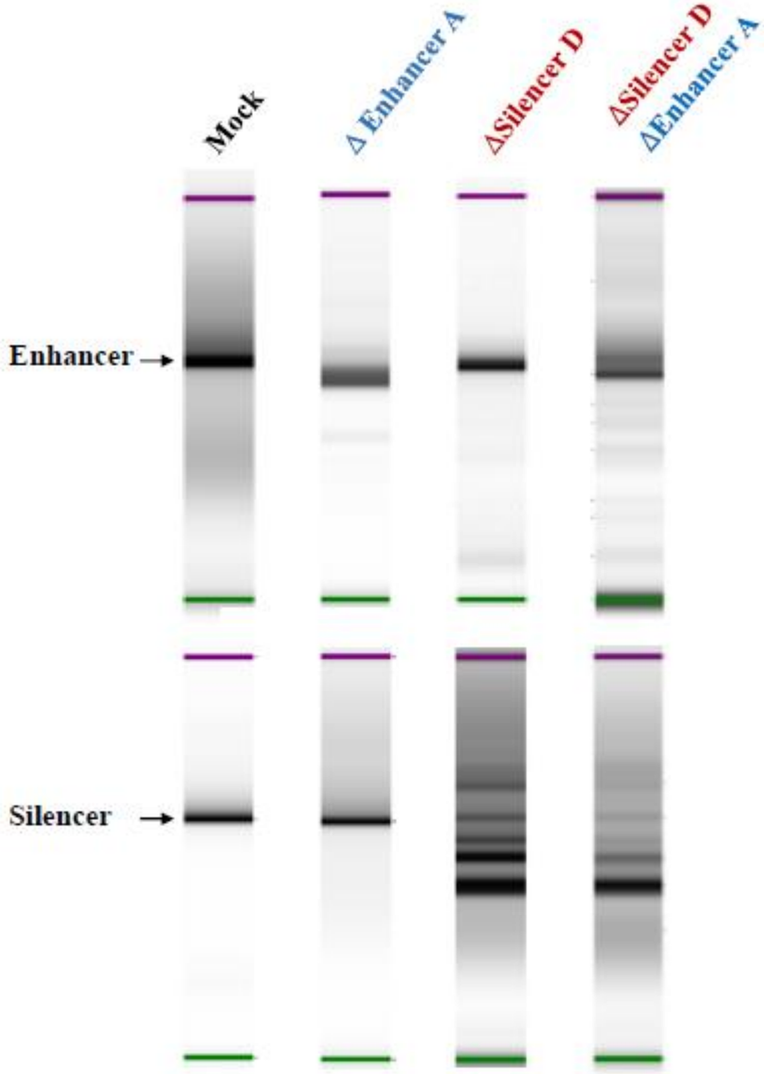


**Supplemental Figure 9. Silencer and enhancer co-deletion**. Electrophoreses of the targeted enhancer and silencer units. Arrows indicting the unit sizes prior to genomic editing.

A

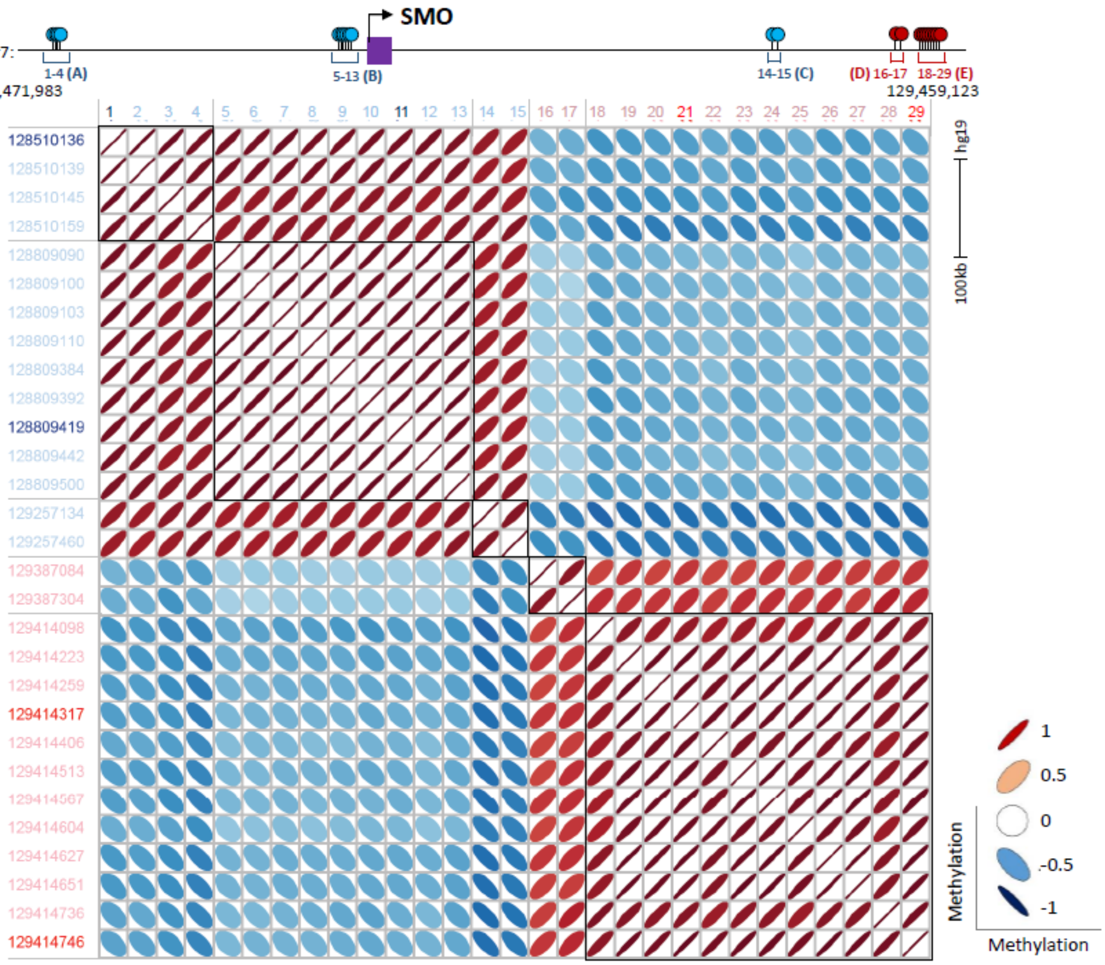

SMO
Chr7:
1-4 (A)
5-13 (B)
14-15 (C)
(D) 16-17
18-29 (E)
128,471,983
129,459,123
hg19
100kb
128510136
128510139
128510145
128510159
128809090
128809100
128809103
128809110
128809384
128809392
128809419
128809442
128809500
129257134
129257460
129387084
129387304
129414098
129414223
129414259
129414317
129414406
129414513
129414567
129414604
129414627
129414651
129414736
129414746
1
0.5
0
-0.5
-1
Methylation
Methylation

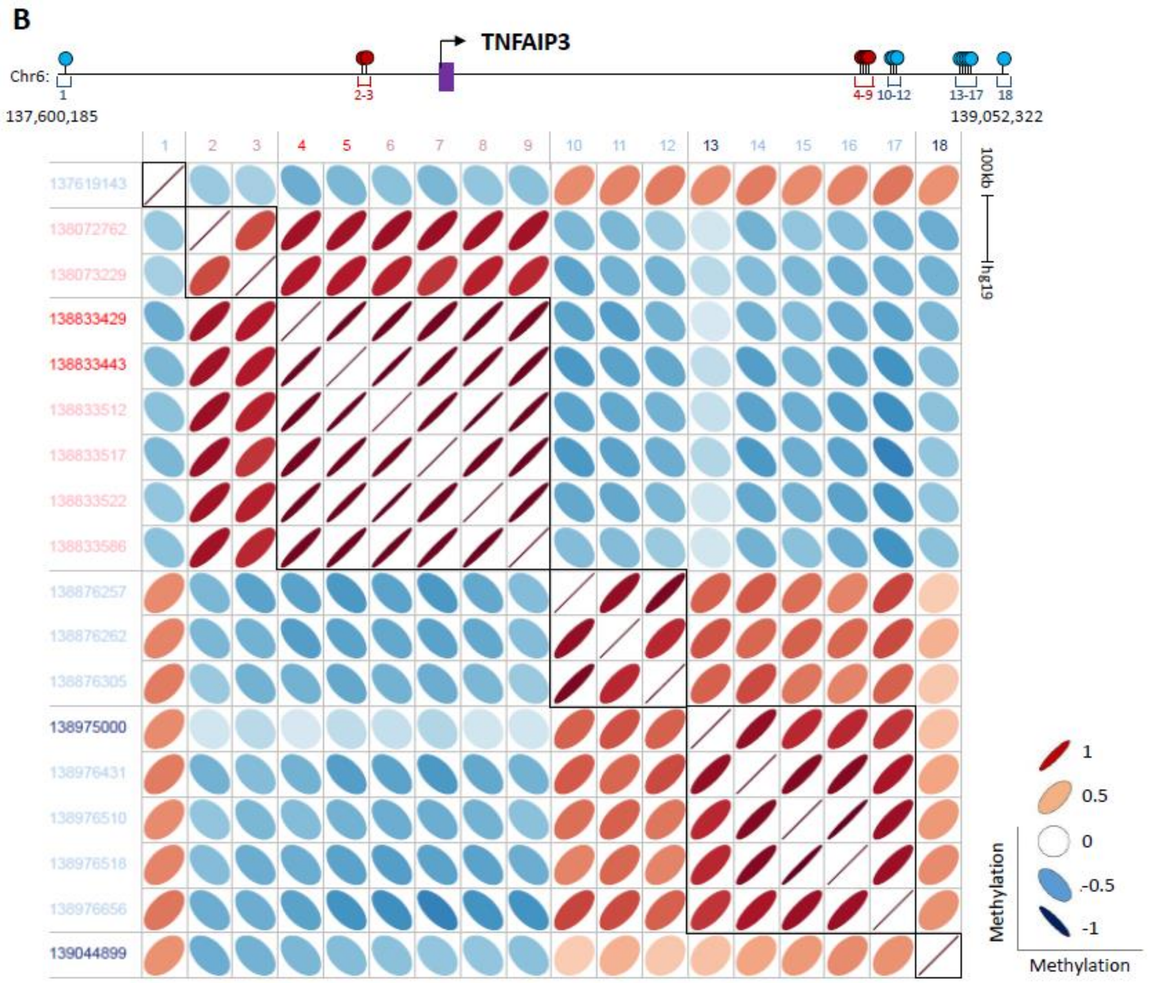


**Supplemental Figure 10. Methylation-methylation coordination maps of genes with multiple regulatory circuits. (A)** Coordination between the methylation levels of SMO-associated sites. Genomic locations of the associated sites are given to the left. **Red label:** positive methylation versus expression associations. **Blue label:** negative methylation versus expression associations. The sites producing best prediction models (see Figure 4) are highlighted. Each square in the matrixes show the methylation versus methylation correlation (R) between two of the associated sites. Genomic maps showing the locations of the associated sites (red bars), of the associated genes (purple), and the site order in the matrix are shown above. **(B)** Same as in A, but for sites associated with the TNFAIP3 genes. The coordination maps of the other analyzed genes are given in Supplemental Data 1

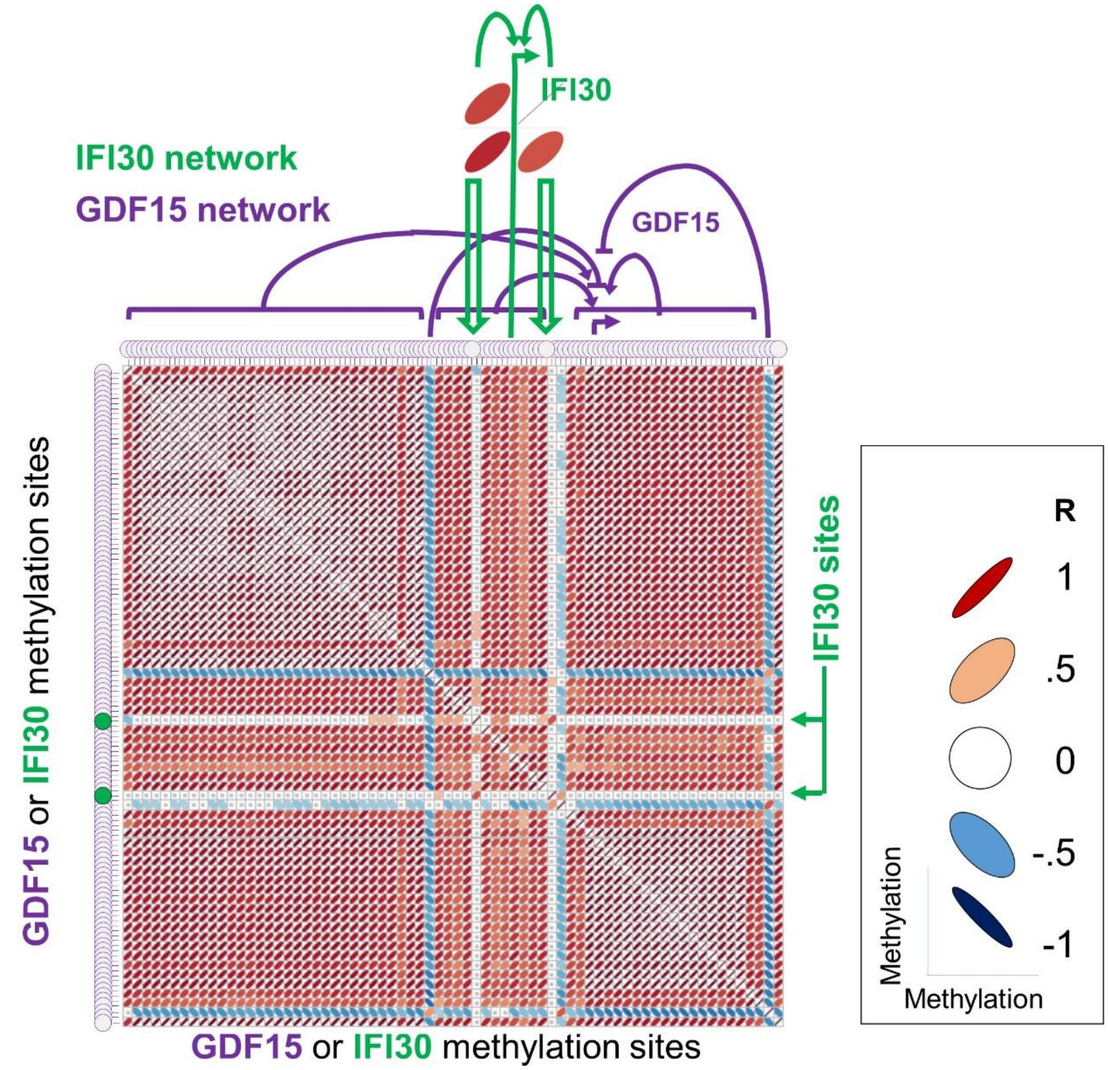


**Supplemental Figure 11. Gene-specific networks**. Matrix showing the coordination between the methylation levels of sites associated with the GDF15 (purple) or the IFI30 genes are shown. Each square in the matrixes show the methylation versus methylation correlation (R) between two of the associated sites. **Blue:** negative methylation versus expression associations. **Red:** positive associations. White squares denote no correlation ($R^2$ <0.1). Maps of the other analyzed overlapping gene domains are given in Supplemental Data 2

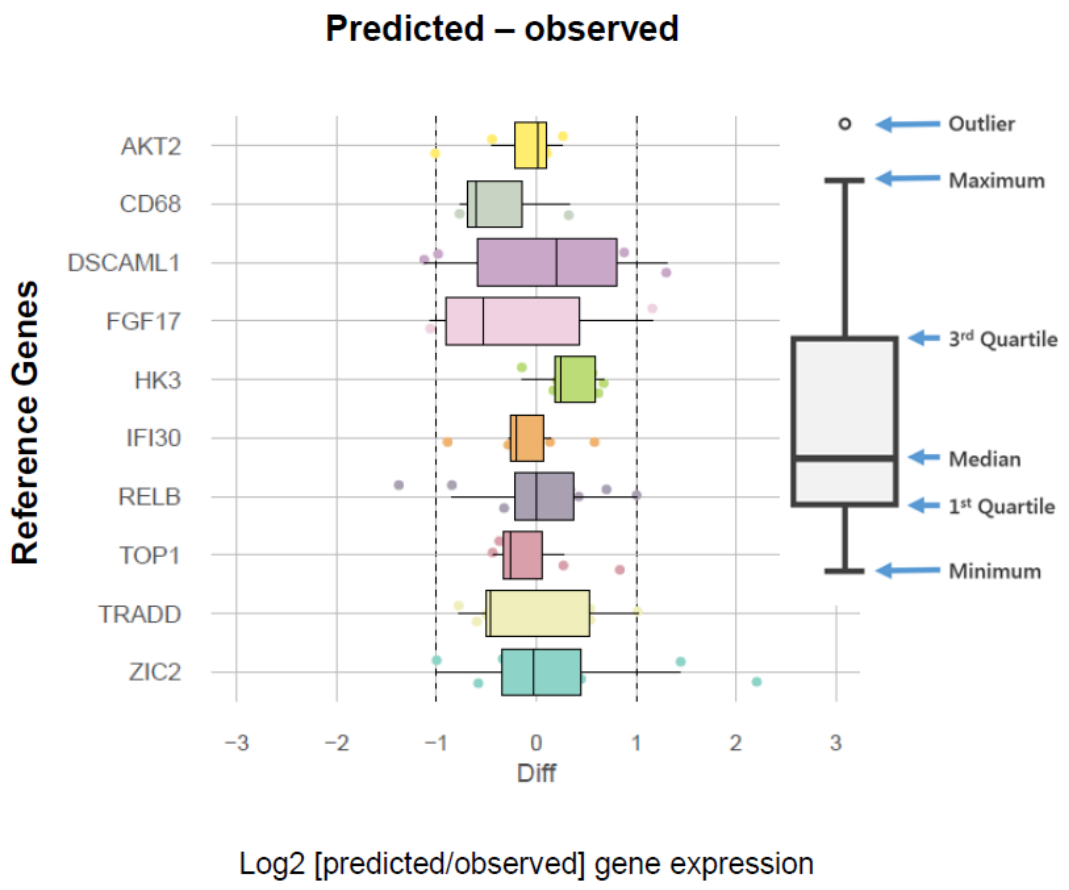


**Supplemental Figure 12.** Log 2 of the differences between observed and predicted gene expression levels for reference (non-driver) genes with developed models. Box plots describing the distributions of prediction accuracy in 24 independent tests.

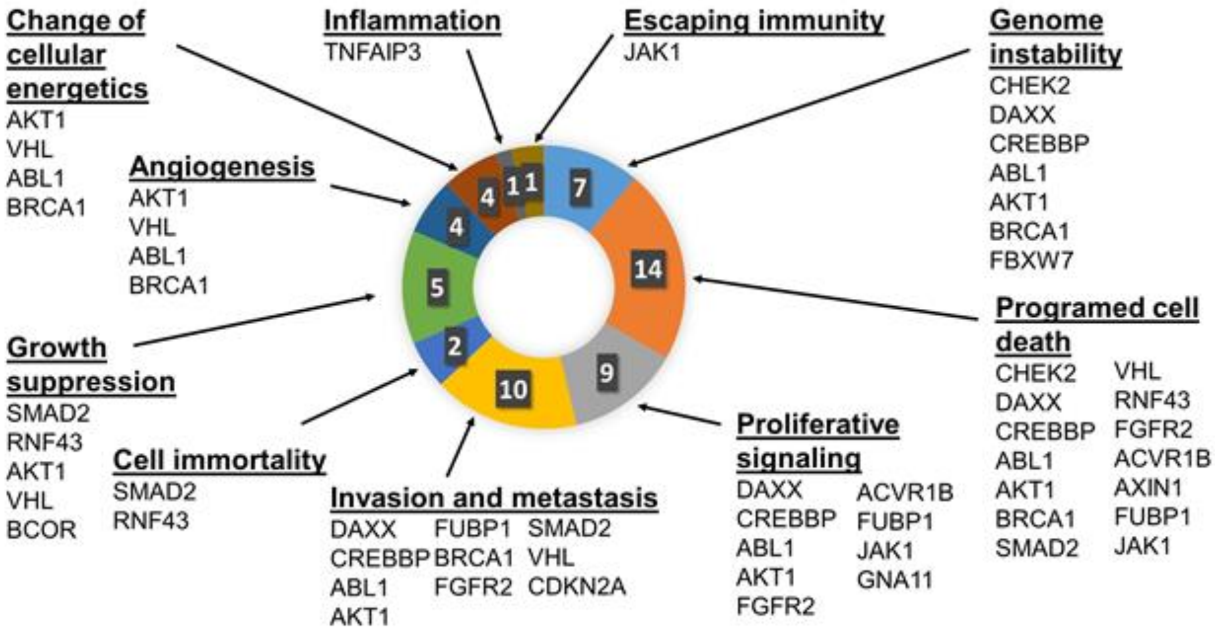


**Supplemental Figure 13.** Cellular functions of misregulated driver genes for which a methylation-based model of expression variation was developed and verified.

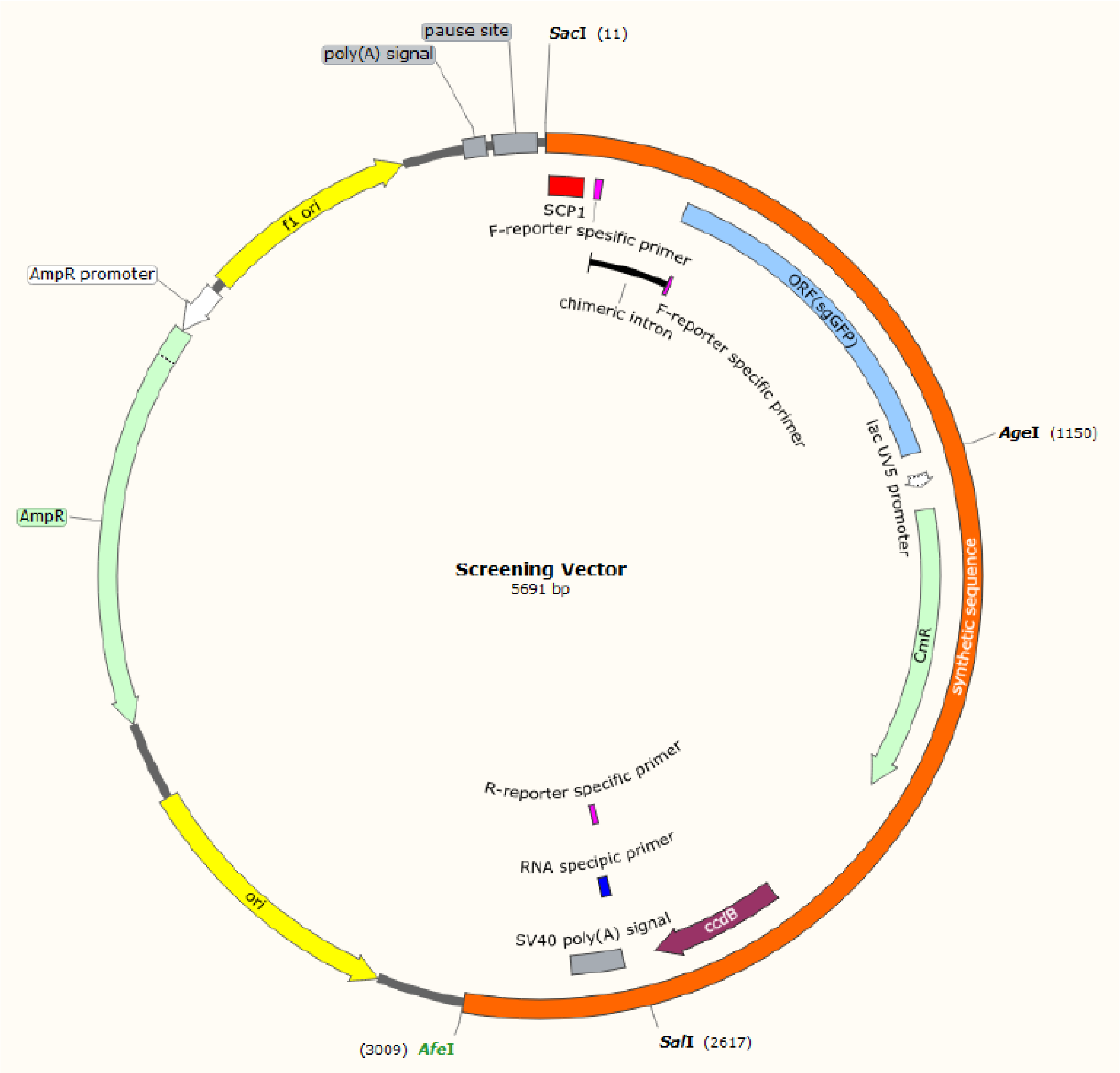


**Supplemental Figure 14. Map of the screening vector used for functional analyses of isolated DNA segments.** The sequence between the SacI and the AfeI sites in the original pGL3-promoter vector (Promega, GenBank accession number U47298) was replaced with the sequences shown here. The modified vector produced a certain amount of basal transcription when no regulatory elements was presented. To evaluate regulatory functionality, putative silencer or enhancer elements were incorporated between the AgeI and the SalI sites.

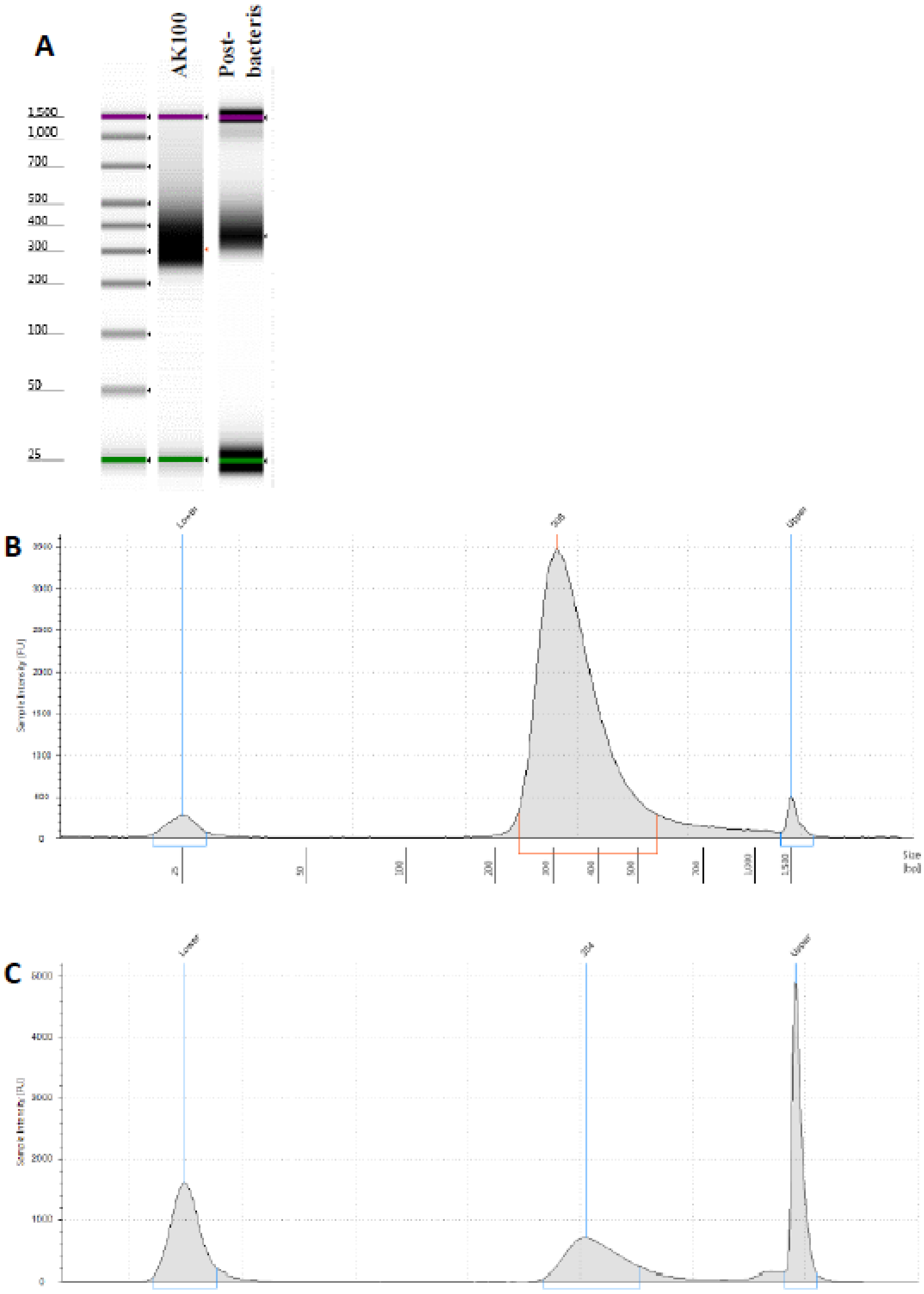


**Supplemental Figure 15.** Gel images **(A)** and size distributions of sample #100 target-enriched library before **(B)** and following **(C)** propagation in bacteria

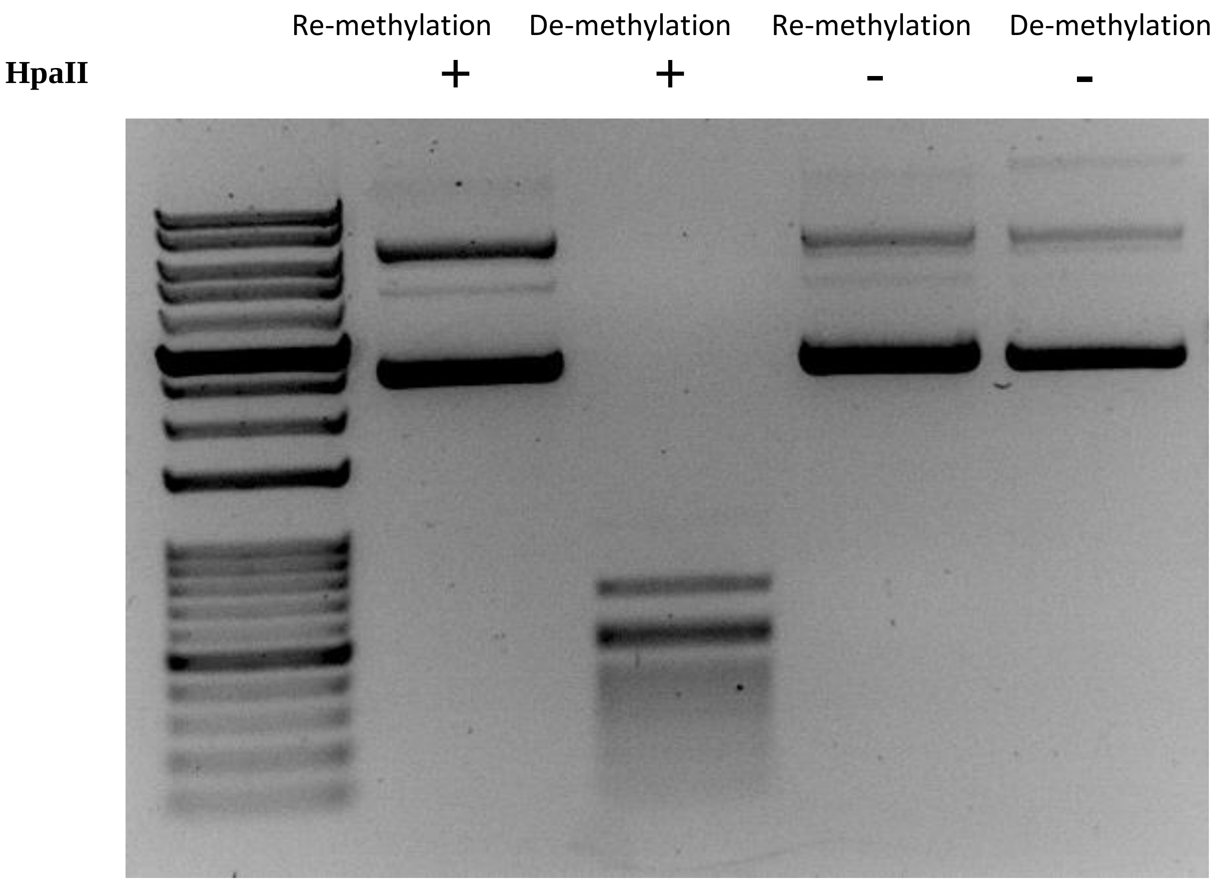


**Supplemental Figure 16.** Efficiency of the methylation assay confirmed by digestion with the methyl-sensetive HpaII restriction enzyme of libraries of captured genomic segments following amplification and propagation in bacteria (de-methylation) or following amplification, propogation in bacteria, and in-vitro methylation (re-methylation).

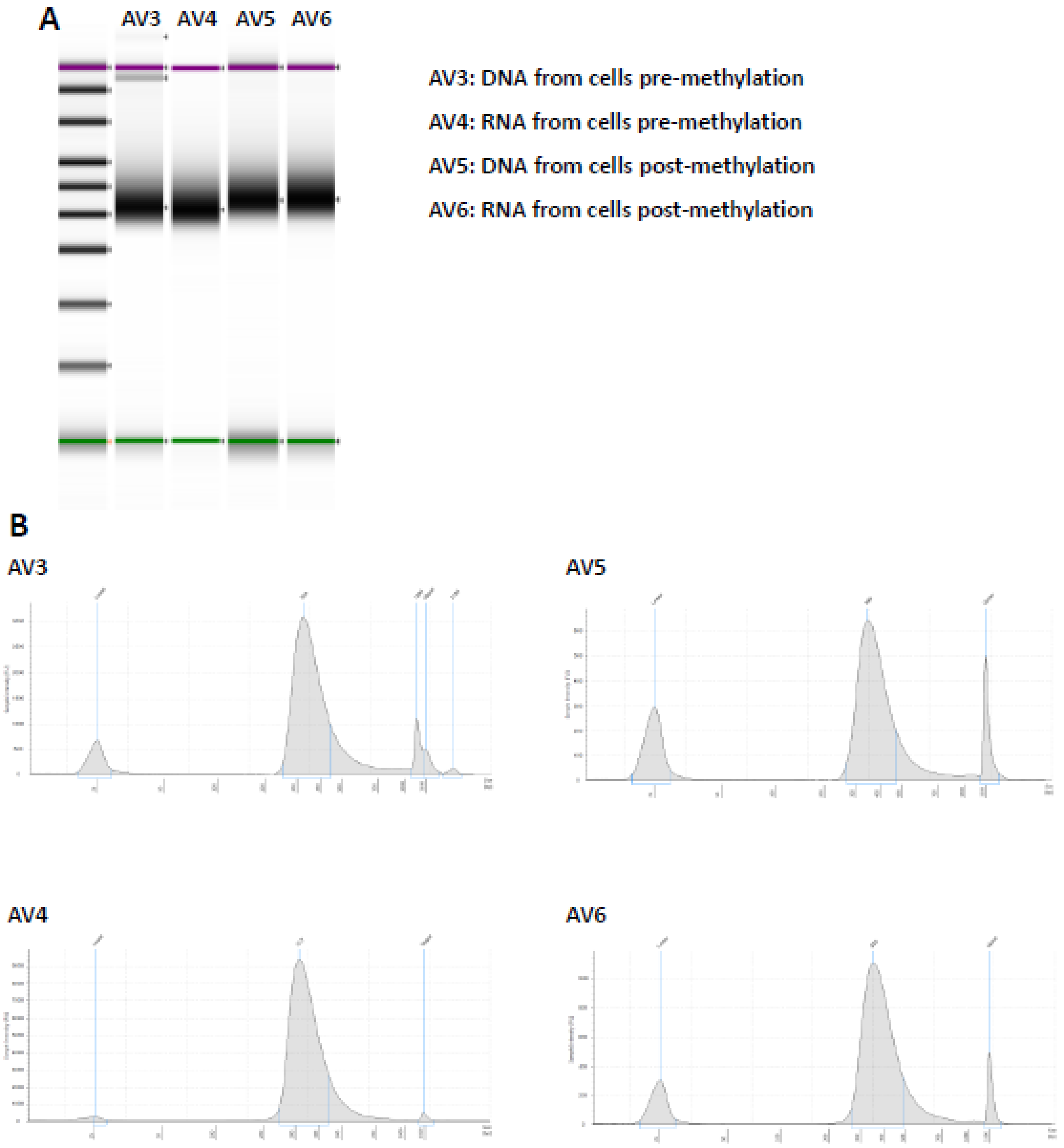


**Supplemental Figure 17.** Gel image **(A)** and size distribution **(B)** of plasmid DNAs and RNAs extracted from T98G cells following transfection with methylated or un-methylated sample #100 libraries.
